# ECD, a novel androgen receptor target promotes prostate cancer tumorigenesis by regulating glycolysis

**DOI:** 10.1101/2025.01.30.635534

**Authors:** Mohsin Raza, Asher Rajkumar Rajan, Benjamin B. Kennedy, Timothy E. Reznicek, Farshid Oruji, Sameer Mirza, M Jordan Rowley, Carsten Stephan, Glen Kristiansen, Kaustubh Datta, Bhopal C. Mohapatra, Hamid Band, Vimla Band

**Author notes:** **Corresponding authors:** Vimla Band, PhD, Department of Genetics Cell Biology & Anatomy, University of Nebraska Medical Center, 985805 Nebraska Medical Center, Omaha, NE, 68198, USA. Hamid Band, MD, PhD, Eppley Institute for Research in Cancer, and Allied Disease, 985950 Nebraska Medical Center, Omaha, NE 68198-5950, USA. Bhopal C. Mohapatra, Department of Genetics, Cell Biology, and Anatomy, University of Nebraska Medical Center 985805 Nebraska Medical Center, Omaha, NE 68198. Both authors contributed equally to this manuscript (Mohsin Raza, Asher Rajkumar Rajan). Department of Chemistry, College of Science, UAEU University, Al Ain, Abu Dhabi, U. A. E. Department of Biochemistry and Molecular Biology, Massy Cancer Center, Virginia Commonwealth University, Richmond, VA, United States.

## Abstract

Androgen receptor (AR)-mediated signaling is essential for PC tumorigenesis. In the TCGA database we observed a positive correlation between ECD and AR expression. Consistently, Dihydrotestosterone (DHT) treatment of PC cell lines increased ECD mRNA and protein levels, and AR knockdown (KD) reduced ECD expression. Bioinformatic analysis predicted three consensus androgen response elements in the ECD promoter, and DHT treatment increased AR occupancy at the ECD promoter, and enhanced ECD promoter activity. Enzalutamide treatment decreased ECD levels, and ECD knockout (KO) in PC cells reduced oncogenic traits, suggesting a functional role of ECD to maintain PC oncogenesis. ECD mRNA and protein are overexpressed in PC patient tissues, and its overexpression predicts shorter survival. Overexpression of ECD in PC cell lines enhanced the oncogenic traits *in vitro* and developed faster and larger highly proliferative xenograft tumors. RNA-seq analysis of mouse tumors revealed an increase in mRNA levels of several glycolytic genes. ECD associates with mRNA of key glycolytic genes and is required for their stability, consistent with our recent demonstration of ECD is an RNA binding protein. Higher glucose uptake and glycolysis was seen upon ECD overexpression in PC cells. Together, we demonstrate the role of a novel AR target gene ECD in PC tumorigenesis.

## Introduction

Prostate cancer (PC) is the second leading cause of cancer related adult males deaths [1]. Androgen receptor (AR) mediated signaling activated through binding of androgens is essential for PC tumorigenesis [2]. Initial androgen deprivation therapy is invariably followed by development of castration resistant PC (CRPC) [3]. AR inhibitors have proven effective but resistance to these inhibitors develops within 2-3 years, even though most such patients retain a functional AR that sustains tumor growth and spread [4, 5]. Thus, understanding of novel molecular pathways that sustain the AR-driven PC tumorigenesis could open novel approaches to therapy of AR-dependent PC.

Most solid tumors, including PC, exhibit metabolic reprogramming to cope with increased macromolecular synthetic, redox and energetic demands for increased oncogenic traits [6]. The hallmark of cancer cells, the Warburg effect, involves a switch to aerobic glycolysis instead of oxidative phosphorylation [7]. Increase in the rate of glycolytic flux has been reported during the late stage of high-grade primary PC, and CRPC, as well as PC with neuroendocrine differentiation [8, 9]. In addition, PC has been shown to maintain a unique androgen regulated metabolic profile with high rates of lipogenesis as well as oxidative phosphorylation [10]. While the role of glycolysis is well accepted across cancers, mechanisms that promote glycolysis in AR-dependent PC remain an area of active investigation. A unique property of prostate epithelial cells is that glycolytic metabolism sustains physiological citrate secretion and in early stages of PC, citrate is needed for oxidative phosphorylation and lipogenesis for tumor progression, however, in advanced and incurable CRPC, a metabolic shift towards choline, amino acid, and glycolytic metabolism takes over tumor progression [11]. Thus, inhibiting glycolysis in advanced PC provides a unique opportunity. Here, we identify ECD as a novel link between AR-dependent PC tumorigenesis by modulation of glycolysis through its novel RNA binding role to stabilize mRNAs of key glycolytic genes.

ECD is the highly conserved mammalian orthologue of *Drosophila* ecdysoneless (ECD) gene [12]. *Ecd* in essential for mouse embryonic development and deletion of *Ecd* in *Ecd^flox/flox^*mouse embryonic fibroblasts (MEFs) led to G1 cell cycle arrest [13]. ECD plays a role in mitigating endoplasmic reticulum stress through upregulation of GRP78 [14]. Previously, we have shown that ECD contains a nuclear export signal and shuttles between the nucleus and cytoplasm [15]. Notably, it interacts with several components of the mRNA spliceosome machinery, and participates in pre-mRNA processing in both drosophila and mammalian cells [16, 17], as well as in mRNA export in mammalian cells [18]. Our [19, 20] and others’ studies have established that ECD mRNA and protein are overexpressed in various solid cancers, and that ECD overexpression (OE) plays a critical role in promoting tumorigenesis [21, 22]. Importantly, transgenic ECD OE targeted to mouse mammary epithelium led to mammary epithelial hyperplasia and tumorigenesis [23]. However, the role and regulation of ECD in PC tumorigenesis has remained unexplored until this study.

Here, we establish that ECD is a novel AR target gene, and its’ expression correlates with AR expression. ECD expression is regulated by AR, through androgen-dependent AR recruitment to the ECD promoter, and androgen treatment increased ECD promoter activity. Enzalutamide treatment decreases ECD levels, as well as ECD KO decreases oncogenic traits, suggesting a key role of ECD in PC oncogenesis. ECD mRNA and protein are overexpressed in PC patient tissues, with ECD OE predicting short survival in PC patients. ECD OE enhanced *in vitro* oncogenic traits, as well as *in vivo* tumorigenicity. Using RNA-seq analysis of xenograft tumors, RNA immunoprecipitation analyses, and functional assays we demonstrate that ECD binds to and stabilizes the mRNAs of key glycolytic genes to promote glycolysis. Thus, our studies identify a novel AR target gene ECD and its function in promoting PC oncogenesis through regulation of glycolysis.

## Results

### ECD expression in prostate cancer cells is regulated by androgen receptor

All PC are initially androgen-driven [24] and most remain dependent on androgen receptor signaling even after castration-resistant [25], Given the connection of AR in PC, we analyzed TCGA-prostate adenocarcinoma dataset and observed a significant positive correlation between AR and ECD mRNA expression **(Fig. 1A)**. To test this *in vitro*, we used AR positive ECD expressing prostate cancer cell lines (**Fig. S1A**): LNCaP (androgen-responsive, derived from a lymph node metastasis of human prostatic adenocarcinoma), C4-2B (an androgen-independent LNCaP subline isolated from bone metastasis in castrated mice [26], 22Rv1 (androgen responsive, human prostate carcinoma cell line [27] and VCaP (androgen sensitive cells derived from vertebral PC metastasis [28]. Despite differing androgen sensitivity, AR expression remains essential for proliferation and survival in these cell lines [29–31]. Consistently, DHT dose-response **(Fig. 1B-E, and Fig. S1B-E)**, as well as time-course analyses (**Fig. 1F-I, and Fig. S1H & I)** in four cell lines LNCaP, C4-2B, 22Rv1, and VCaP cell lines showed ECD protein and mRNA levels increased rapidly, plateaued at later time points and decreased towards the end of the time course. Decreased at later time points is similar to other AR target genes, such as PSA, possibly due to limited DHT due to its metabolism over longer time in charcoal-stripped serum [32]. Further, the basal as well as DHT-induced increase in ECD protein levels were abrogated by siRNA KD of AR, which was confirmed by anti-AR immunoblotting in LNCaP **(Fig. 1J)**, C4-2B **(Fig. 1K),** 22Rv1 **(Fig. S1J),** and VCaP **(Fig. S1K)** cells with or without the addition of exogenous DHT. These results show that ECD expression in PC cell models is positively regulated by androgens and dependent on the AR, suggesting the likelihood that ECD is transcriptionally regulated by AR.

**Fig. 1.**
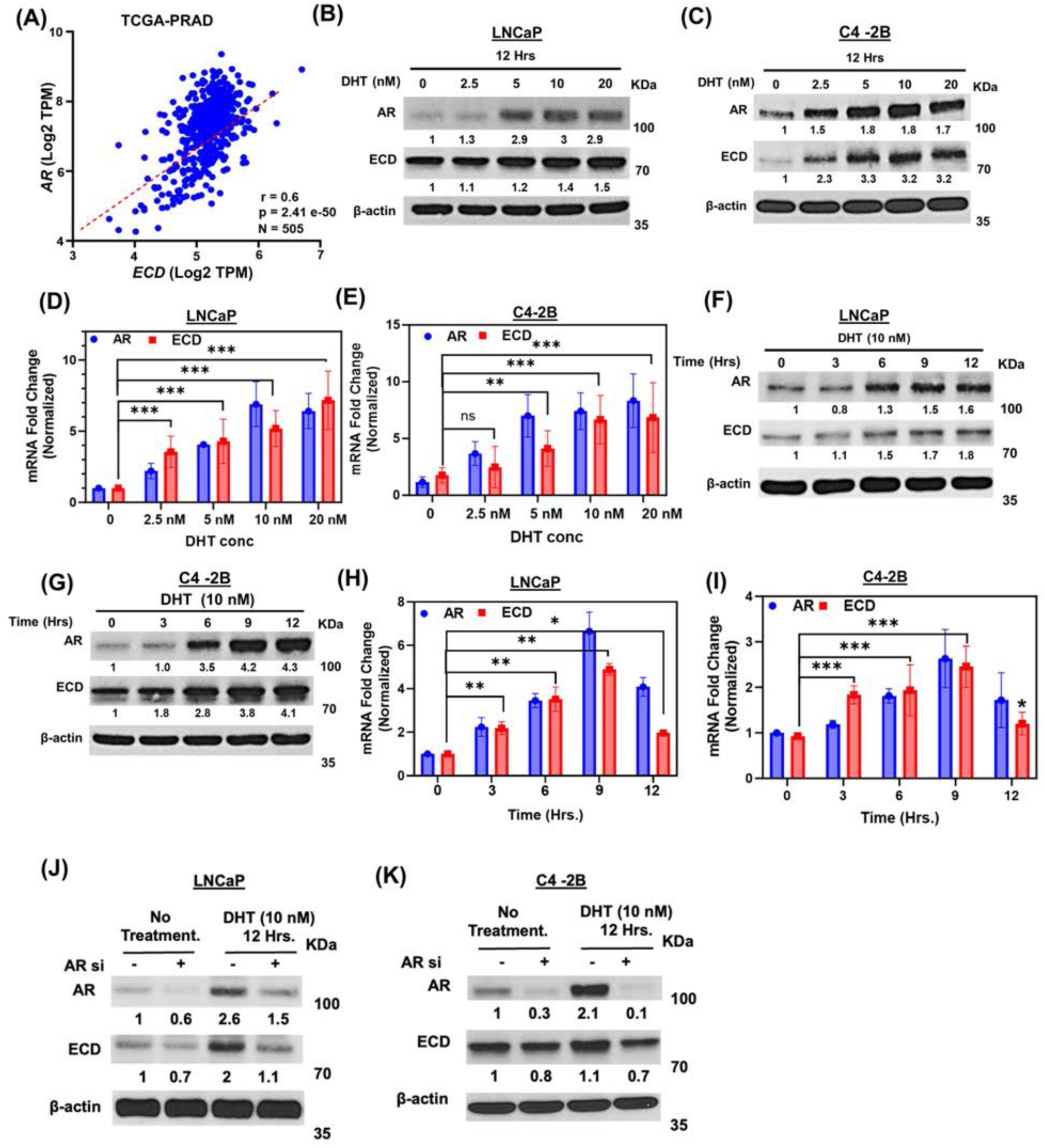
Correlation of ECD and AR expression; AR regulates ECD expression. **(A)** Spearman’s pairwise correlation plot illustrates a positive correction of AR and ECD mRNA in TCGA Prostate Adenocarcinoma Firehose dataset of cBioPortal online platform. Number of patients (N), correlation coefficient (r) and p-value are indicated on the plot. Pairwise correlation analysis was performed to calculate statistical significance. **(B & C)** Indicated PC cell lines were cultured in steroid free conditions for 72 hours. Cells were then treated with indicated concentrations of DHT for 12 hours and immunoblotted with anti-ECD or anti-AR antibodies. β-actin was used as a loading control. **(D & E)** qRT-PCR was performed in the same samples; RNA was isolated by standard TRIzol phenol chloroform method. 18s rRNA was used for normalization. **(F-I)** Similar protocols as mentioned above were used to perform a time response with DHT. Western blotting **(F & G)** and qRT-PCR **(H & I)** were performed on cell lysates, as above. **(J & K)** Indicated PC cell lines transfected with control siRNA or AR siRNA and cultured for 72 hours. Cells were then treated with indicated concentrations of DHT for 12 hours and then lysates were western blotted with anti-ECD or anti-AR antibodies. β-actin was used as a loading control in all blots. Numbers below the blots show the quantification of band intensities after normalizing with their respective loading control, β-actin in comparison with control samples using ImageJ software. Bar graphs represent mean ± SEM from three independent experiments, each done in triplicates. Student t- test was used to calculate statistical significance *** p < 0.001, ** p < 0.01, * p < 0.05, ns p > 0.05.

### AR binds to the ECD promoter to regulate ECD expression, and Enzalutamide treatment decreases ECD expression

Both androgens activated and constitutively active ARs function by binding to gene promotors through defined androgen response elements (AREs) to regulate transcription of downstream target genes [33]. Given that DHT induction, combined with AR dependence of *ECD* mRNA expression, we examined the ECD promoter region for potential AR binding sites, using three independent prediction tools, Pscan [34], EPD motif search tool [35] and de novo motif discovery algorithm Motif Based Sequence analysis tool [36]. These analyses revealed three potential androgen response elements (AREs) in the ECD promoter: nucleotides −698 to - 682 (site 1); −657 to −641 (site 2); and −640 to −624 (site 3) **(Fig. 2A & B)**. Indeed, chromatin immunoprecipitation (ChIP) assays demonstrated specific occupancy of AR at the ECD promoter (−712 to −565, encompassing all three sites) in all four, LNCaP, C4-2B, 22Rv1, and VCaP PC cell lines compared to IgG control, with the occupancy significantly increased upon DHT treatment **(Fig. 2C & D, and Fig. S2A & B)**. Next, we cloned a ∼1,500 bp region encompassing the ECD promotor region in a reporter construct upstream of secreted Gaussia Luciferase. PC cells transduced with the ECD promotor Luciferase construct were left untreated or treated with DHT for 3 hours, and secreted Gaussia Luciferase and alkaline phosphatase (SEAP; internal control) activities were measured in the media. Notably, compared to basal reporter activity (Ctrl; first bar), DHT treated cells expressing the Full length ECD promoter showed about 2.5-fold increase in luciferase activity **(Fig. 2E & F, lane 2)**. Importantly, constructs with deletion of AREs either alone or in combination showed progressive diminution of luciferase activity while the construct with all three AREs deleted showed activity close to that of the control **(Fig. 2E & F, lanes 3-9).** Collectively, our results identify ECD is an AR-regulated gene, with three functional ARE elements in its promoter region that mediate androgen-induced AR-dependent ECD expression.

**Fig. 2.**
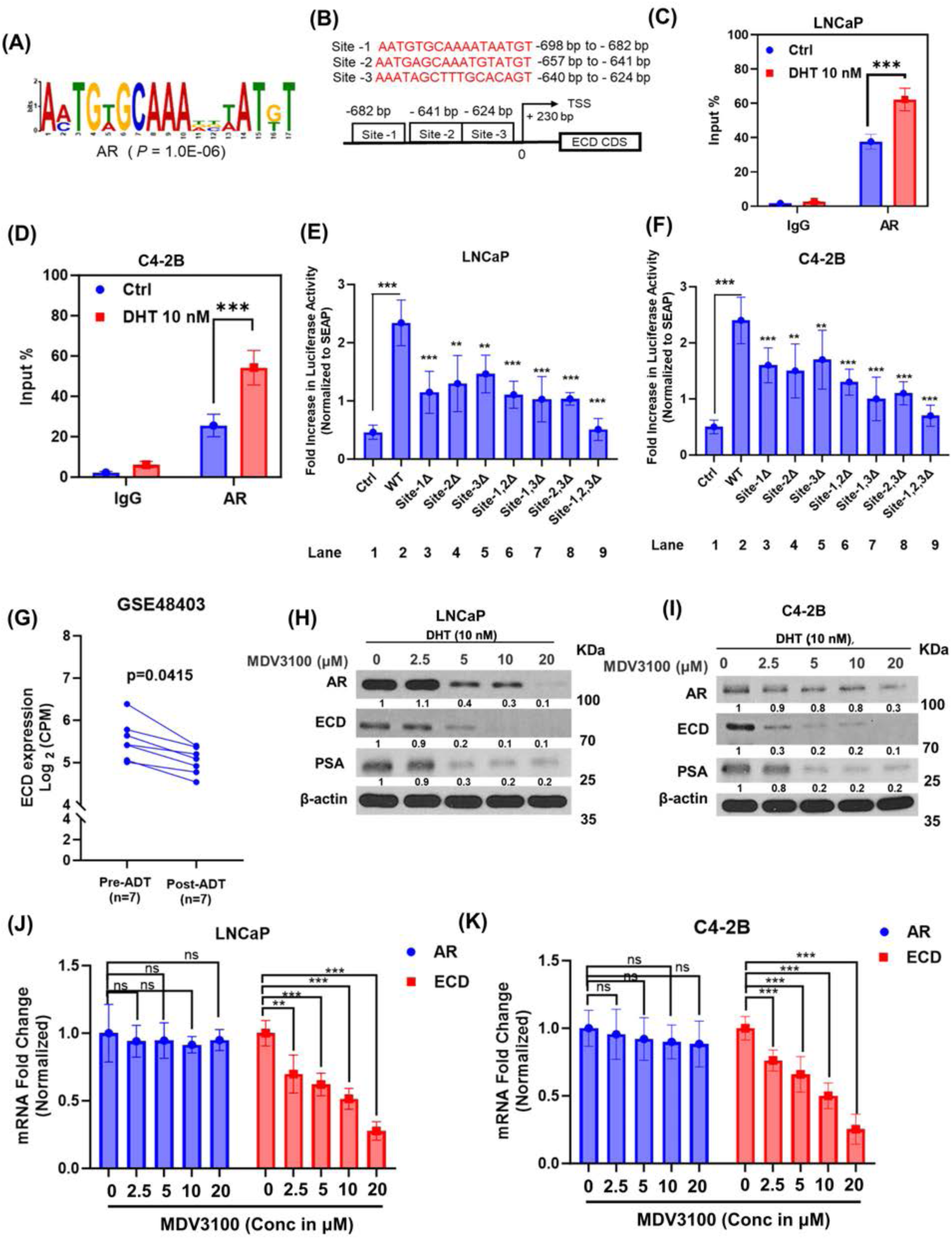
Androgen treatment increased AR occupancy on ECD promoter and Enzalutamide treatment decreases ECD expression. **(A & B)** AR binding site; Top three predicted AR binding sites on the ECD promotor. **(C & D)** Chromatin immunoprecipitation (ChIP) analysis. Indicated cells were cultured in steroid-free conditions for 3 days, followed by dihydrotestosterone (DHT) treatment for 3 hours. Cells were cross-linked with formaldehyde and chromatin was immunoprecipitated with antibody against AR. IgG, used as a non-binding Control. Immunoprecipitated DNA was quantified with qPCR for specific binding sites at ECD promoter using primers encompassing all three sites −712 to −565 from transcription start site. Student t- test was performed to calculate statistical significance *** p < 0.001. **(E & F)** DHT increases ECD promoter activity. ECD promoter spanning approx. 1500 bp was cloned upstream of secreted Gaussia Luciferase, Secreted Alkaline phosphatase (SEAP) was used as an internal control. Cells expressing WT promoter or various AR binding sites deleted mutants were treated with 100 nM DHT for 3 hours. Supernatant was collected, and Luminescence was measured using Secrete-Pair Dual Luminescence Assay Kit. Data represents three independent experiments, each done in triplicates. *Student t- test* was performed to calculate statistical significance *** p < 0.001, ** p < 0.01. **(G)** Box plot depicting ECD mRNA expression in prostate tumors pre- and post-androgen deprivation therapy 7 samples in each case (GSE48403). A Wilcoxon matched-pairs signed-rank test was used to compare expressions in the groups. **(H & I**) Indicated PC cell lines were cultured in steroid-free medium for 72 hours, cells were then treated with 10 nM DHT, followed by treatment with indicated concentration of MDV3100 for another 48 hours. Lysates were collected and western blotted with indicated antibodies. β-actin was used as a loading control. **(J & K)** qRT-PCR was performed in the same samples where the RNA was isolated by standard TRIzol phenol chloroform method. 18s rRNA was used for normalization. Bar graphs represent fold change in AR and ECD mRNA in LNCaP **(J)** and C4-2B cells **(K)** respect to vehicle treatment. Data represent mean ± SEM from three independent experiments, each done in triplicates. *Student t- test* was performed to calculate statistical significance *** p < 0.001, ** p < 0.01.

Based on a published report GSE 48403 [37] showed ECD mRNA expression was decreased post-androgen-deprivation therapy as compared to pre-treatment **(Fig. 2G),** we treated LNCaP, C4-2B, 22Rv1, and VCaP cell lines with different concentrations of Enzalutamide in the presence of DHT. Notably, Enzalutamide treatment dose-dependently decreased ECD protein levels in all four cell lines **(Fig. 2H & I, and Fig. S2C & D)**. Enzalutamide is reported to inhibit AR binding to androgen and destabilize AR by dissociating it from the HSP90 complex, resulting in reduced AR protein stability [38, 39], consistent with this, we observed a decrease in AR and PSA, an AR target gene **(Fig. 2H & I, and Fig. S2C & D)**. Enzalutamide induced reduction in ECD is at the transcriptional levels **(Fig. 2J & K, and Fig. S2E & F)**. Together, these results conclusively demonstrate ECD is a novel AR target gene.

### ECD expression is required to sustain oncogenic traits of prostate cancer cells

Next, we engineered the androgen-dependent LNCaP and VCaP cell lines, as well as androgen-independent C4-2B and 22Rv1 cell lines for doxycycline (Dox)-inducible CRISPR knockout with two distinct guide RNAs (gRNA1 and gRNA2) against ECD. Dox-inducible depletion of ECD in gRNA1/2 expressing cell lines compared to the non-targeted control (Cas9-NTC) was observed **(Fig. 3A, and Fig. S3A & B).** Analysis of colony forming capacity as a measure of cell growth demonstrated that ECD depletion led a significant decrease in PC cell proliferation compared to NTC **(Fig. 3B & C)**. Further, ECD depletion in comparison with NTC exhibited a significant decrease in oncogenic traits, such as tumorsphere forming ability **(Fig. 3D & E, and Fig. S3C & D),** a measurement of the stem cell ability of tumor cell lines [40], anchorage independent growth using the soft agar colony forming assay **(Fig. 3F & G, and Fig. S3E & F)**, three-dimensional growth in Matrigel **(Fig. 3H & I, and Fig. S3G & H),** cell migration **(Fig. 3J & K, and Fig. S3I & J),** as well as tumor cell invasion **(Fig. 3L & M, and Fig. S3K & L).** Given that AR is a primary driver of prostate carcinogenesis, and ECD being a transcriptional target of AR, we investigated the extent to which AR-mediated induction of ECD contributes to AR-driven oncogenic processes. To address this, we performed siRNA-mediated knockdown of AR and rescue with ECD in two PC cell lines LNCaP and C4-2B (**Fig. S4A & B)**. Following knockdown and rescue, we conducted soft agar colony formation and Matrigel organoid formation assays to assess oncogenic potential. These experiments revealed that overexpression of ECD partially mitigated the suppressive effects of AR knockdown on soft agar colony (**Fig. S4C-E)** and organoid formation (**Fig. S4F-H)**. These results highlight ECD as a potential downstream target of AR signaling in prostate cancer. Taken together, these results establish a critical role of ECD expression in sustaining the oncogenic traits of PC cells.

**Fig. 3.**
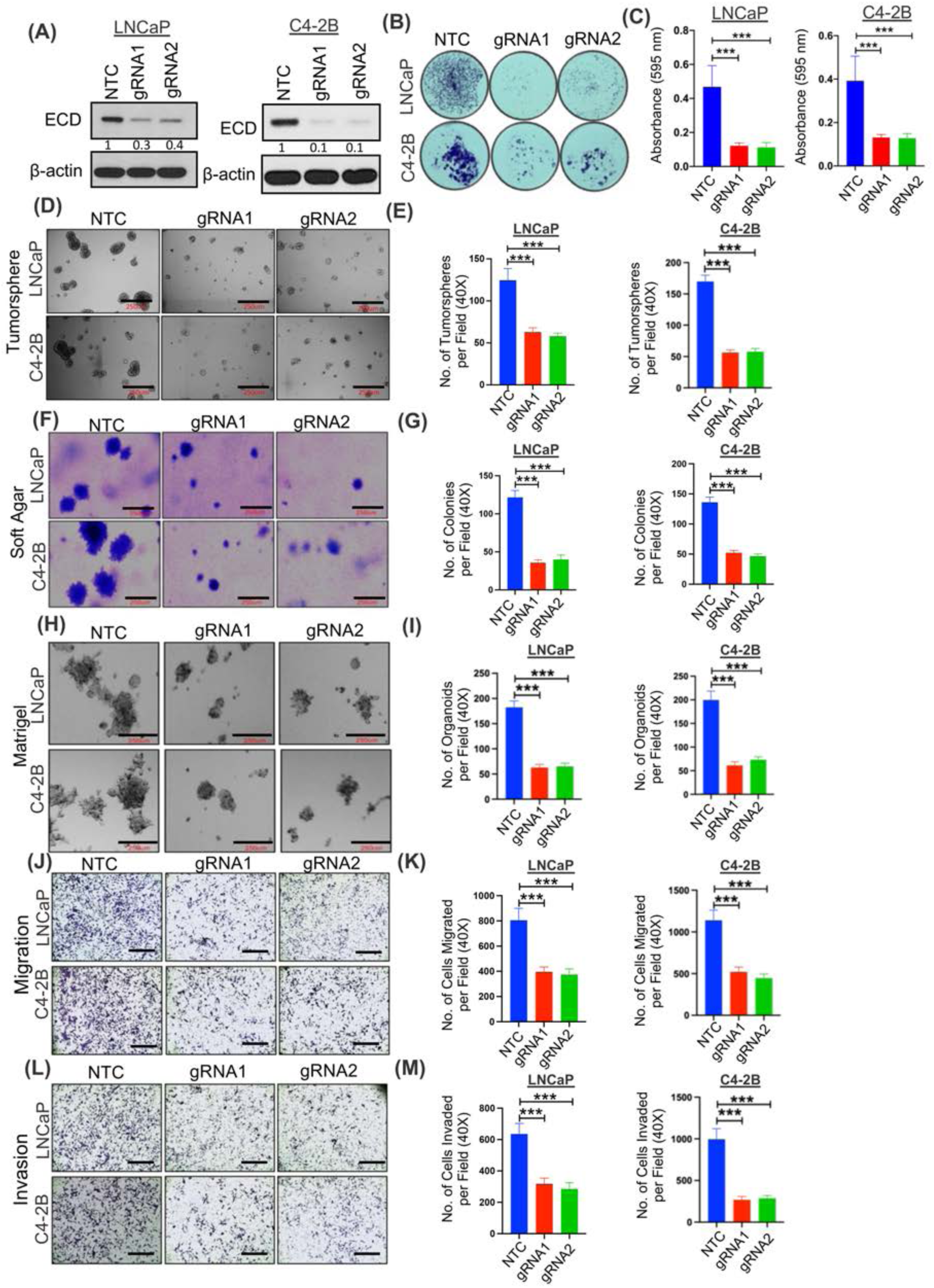
Depletion of ECD induces proliferation block and diminishes oncogenic traits in prostate cancer cells. LNCaP and C4-2B cells expressing doxycycline (Dox) regulated Cas9 and guide RNAs targeting either ECD (gRNA1 & 2) or non-targeting control (NTC) were cultured in the presence of Dox (1 µg/ml) for 96 hours to deplete ECD. **(A)** Western blot analysis. Numbers below the blots show the quantification of band intensities after normalizing with their respective loading control, β-actin in comparison with NTC using ImageJ software. Representative images and histograms depicting 2D (two dimensional) colony **(B & C),** tumorsphere **(D & E**), soft agar colony **(F & G),** Matrigel organoid formation **(H & I),** migration **(J & K)** and invasion **(L & M),** of control and ECD depleted PC cells lines. The representative images of tumorspheres, soft agar colonies and Matrigel organoids are 200x magnification, scale bar, 250 µm. Migration and invasion images are 40x magnification, scale bar, 1000 µm. The Bar graphs represent mean values ± SEM of three experiments done in triplicates from 40x magnification images. *Student t- test* was used to calculate statistical significance *** p < 0.001, ** p < 0.01.

### ECD protein is overexpressed in prostate cancer tumor tissues, and its overexpression predicts shorter patient survival

To assess the expression levels of ECD in primary PC tumor samples, we carried out IHC analysis of a PC tissue microarray (TMAs) comprising of 522 patient samples with available clinical, pathological and survival data using a previously validated anti-ECD antibody [13]. PC samples showed a range of ECD IHC staining. The intensity of staining was expressed as a semiquantitative scoring system: negative = 0, low = 1, moderate = 2 and High =3. Frequency and percentage of patients in each score were assigned and listed (**Fig. 4A**). Based on the intensity of staining, the patients were grouped into ECD-low (staining intensity of 0 or 1) and ECD-high (staining intensity of 2 or 3) groups; representative examples of low IHC staining of hyperplastic tissue, and PC tumor samples low vs. high ECD IHC staining are presented **(Fig. 4B & C)**. Overall, ECD protein was overexpressed in tumor samples of ∼40% of PC patients **(Fig. 4D)**. Kaplan Meier analysis showed that OE of ECD protein correlated with shorter overall survival (p=0.024) **(Fig. 4D)**. The case processing summary table describes the number of events where patient outcome was available, censored are the patients where outcome was unknown, and number of patients used for survival analysis with ECD-low (0&1 score) and ECD-high (2&3 score) (**Fig. 4E)**. Multivariate survival analysis was performed using Cox regression model to evaluate the influence of ECD expression, tumor pT stage, Gleason grade, R-status, age, and pre-operative PSA, on a patient’s overall survival. Patients’ overall survival was significantly dependent on ECD expression (relative risk, RR= 1.69, p= 0.02), tumor pT stage (RR= 3.12, p= 0.0), Gleason grade (RR=2.4, p=0.009), and R-status (RR=1.66, p=0.03 whereas age and pre-operative PSA failed significance **(Table S1).** This analysis independently confirmed the high expression of ECD as a poor prognostic marker, with a significantly elevated relative risk. In addition, analysis of two publicly available mRNA expression datasets (GSE32269 & GSE35988) revealed *ECD* mRNA overexpression in PC tissues as compared to normal prostate **(Fig. S5A & B).** Kaplan-Meier analysis of The Cancer Genome Atlas (TCGA) dataset of 484 mixed prostate adenocarcinoma samples (2022-v32) using the R2 Genomics platform showed that high ECD mRNA was associated with shorter patient survival **(Fig. 4F)**. Collectively, our results show that ECD mRNA and protein levels are increased in a significant subset of PC tumor samples, and that ECD OE correlates with negative prognostic features and shorter survival of patients, suggesting that ECD OE may promote PC tumorigenesis.

**Fig. 4.**
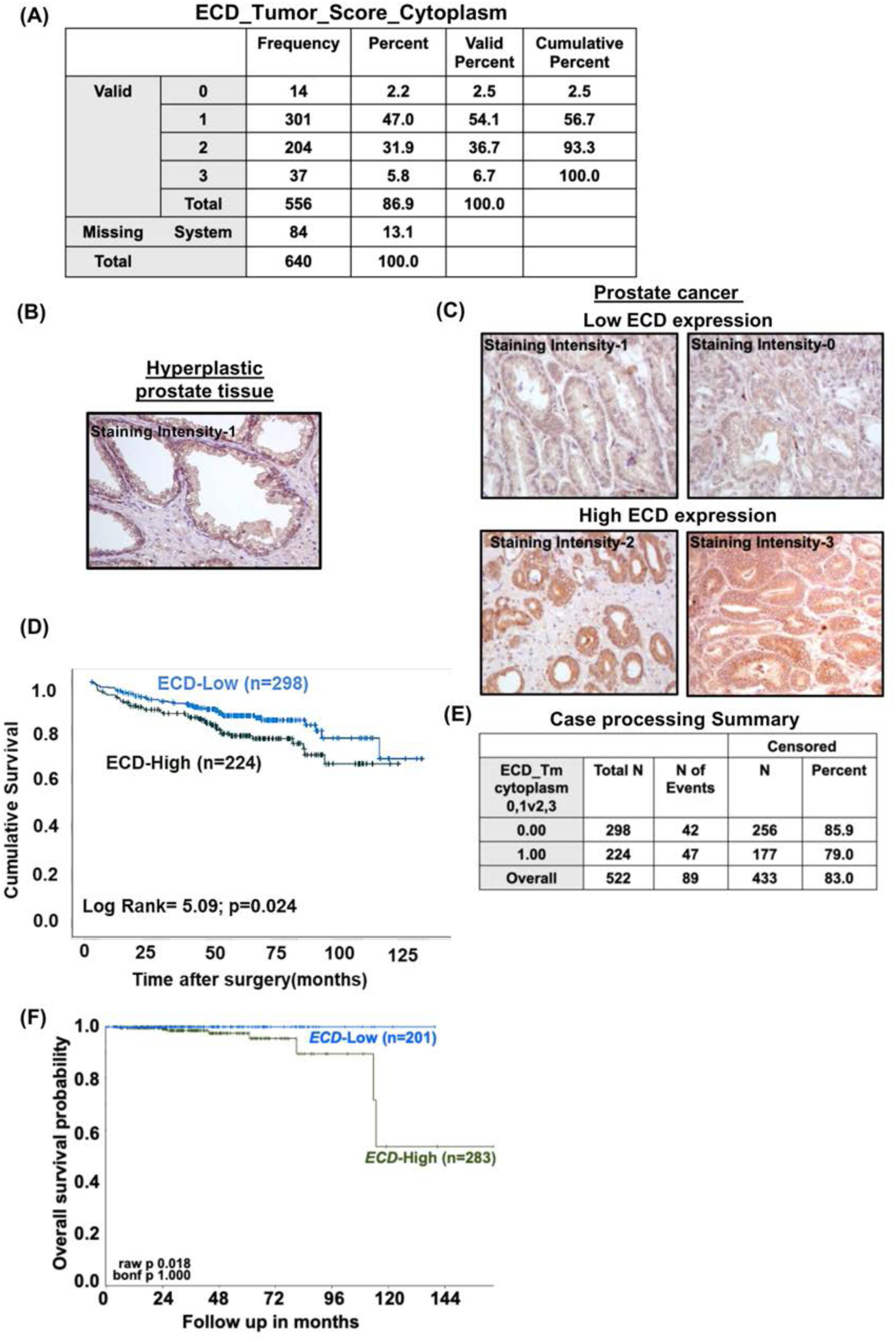
Higher expression of ECD protein and mRNA in prostate cancer tissues correlate with short patient survival. **(A**) Frequency and percentage of patient TMA samples assigned to each staining intensity. **(B & C)** Representative IHC analysis of ECD expression in hyperplastic prostate tissue **(B),** and low-ECD (staining intensity 0, 1) and high-ECD expression (staining intensity 2, 3) in prostate cancer tissues **(C)** at magnification 200x. **(D)** Kaplan-Meier cumulative survival curve after surgery with patient tumors expressing low (staining intensities 0 or 1) or high ECD protein levels (staining intensities 2 or 3). P-values were calculated with a log-rank test. **(E)** Case processing summary with number and percent of events used for the overall survival analysis in Panel D is enlisted. N= number of patients, N of events= number of patients where outcome was known and censored means where the patients’ outcome was unknown **(F)** Kaplan-Meier plot for survival curve of prostate cancer patients shows short survival in patients with tumors expressing high *ECD* mRNA (n=283) in comparison to low *ECD* mRNA (n=201). P-value is indicated in the KM plot. Mixed prostate adenocarcinoma (2022-v32) cohort (n=484) of The Cancer Genome Atlas (TCGA) dataset was used to stratify *ECD* mRNA expression as high (> 31.49 transcripts per million (TPM) and low (< 31.49 TPM). Analysis was performed using R2 Genomics platform.

### ECD overexpression in PC cells promotes the *in vitro* oncogenic traits and *in vivo* tumorigenesis

Next, we established stable ECD-OE PC cell lines LNCaP (hormone sensitive), its derivatives C4-2B (hormone insensitive), and 22Rv1 (hormone insensitive). Overexpression of ECD is observed in all three LNCaP **(Fig. 5A),** C4-2B **(Fig. S6A),** and 22Rv1 **(Fig. S6B)** cell lines. While there were no significant differences in proliferation of vector vs ECD-OE PC cell lines when measured in two-dimensional culture **(Fig. S6C-E),** a significant increase in anchorage independent growth in soft agar **(Fig. 5B & C, and Fig. S6F-I)**, as well as in three-dimensional Matrigel growth **(Fig. 5D & E, and Fig. S6J-M)** was seen in three (LNCaP, C4-2B, and 22Rv1) ECD-overexpressing vs. vector control cells. Notably, ECD-OE LNCaP cells remain hormone sensitive like control cells **(Fig. S6N).**

**Fig. 5.**
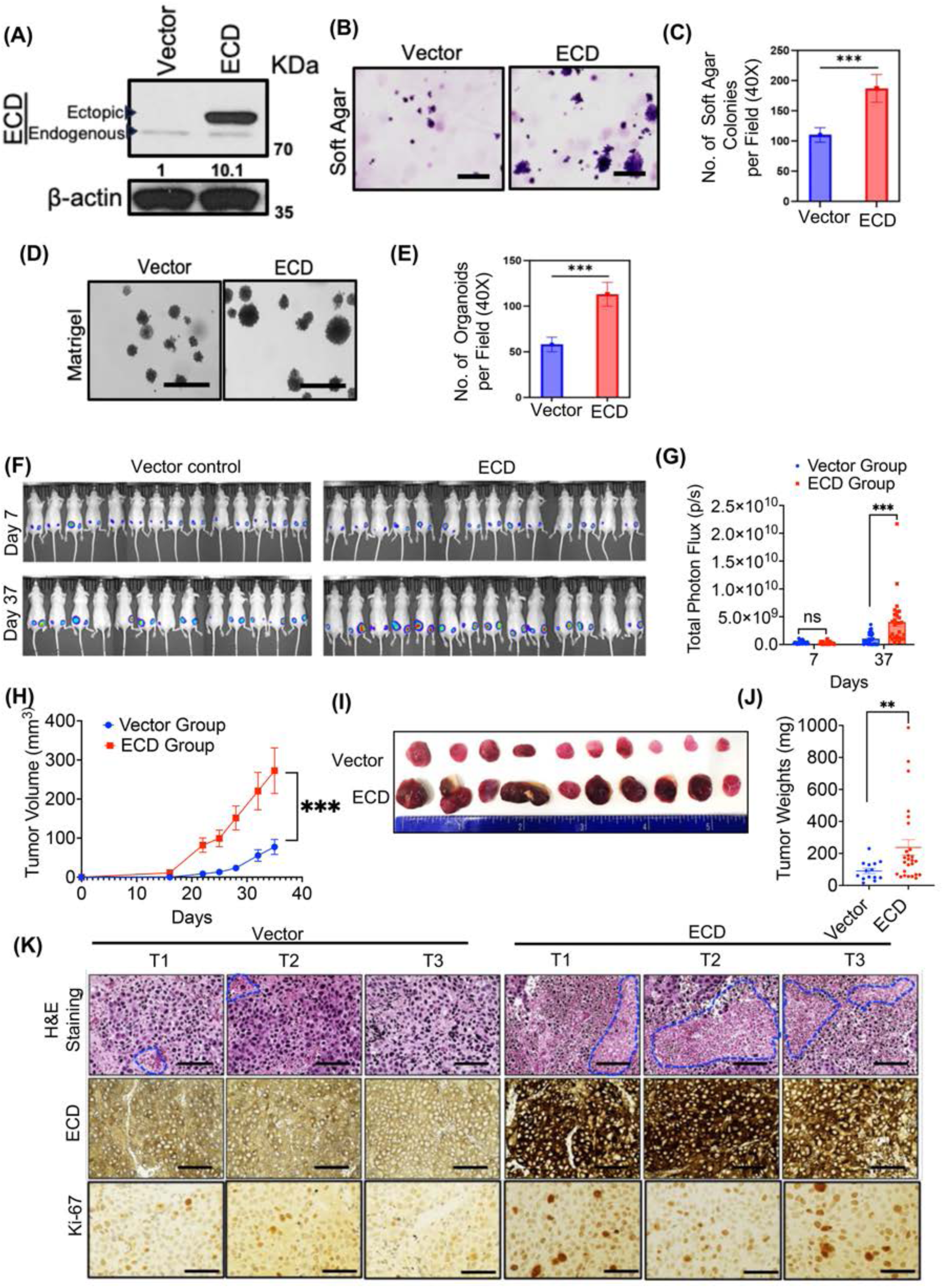
ECD overexpression in LNCaP cells exhibit enhanced *in vitro* oncogenic traits and *in vivo* tumorigenesis. **(A)** Western blot depicts ECD overexpression; densitometry in respect to vector cells after normalizing with β-actin, shown below. **(B)** Representative images in 100x magnification (scale bar, 400 µm) show soft agar colony formation abilities of cells. **(C)** Bars depict the quantitation of soft agar colonies per field at 40x magnification. **(D)** Representative images in 100x magnification (scale bar, 400 µm) depict 3D colony formation abilities in Matrigel. **(E)** Bar graphs show the quantitation of 3D colonies per field at 40x magnification. Data represents three independent experiments, each done in triplicates. ***p < 0.001, Student t-test was used to calculate statistical significance in (C) and (E)**. (F)** mCherry-luciferase expressing LNCaP-vector and ECD-OE cells were injected subcutaneously into nude mice (14/group) in both sides. Primary tumor growth was monitored by bioluminescence imaging at the indicated time on day 7 and day 37. **(G)** Bioluminescence signals of tumors over time are shown as change in photon flux over time. Data are mean ± SEM, n=14 (two sided tumors); Unpaired t-test; ***p < 0.001. **(H)** Tumor growth is followed over days with vernier caliper and fold change in tumor volume is plotted. Data are mean ± SEM, n=14; Two-way ANOVA (mixed model); ***p < 0.001. **(I)** Ten representative harvested tumors are displayed in each group. **(J)** Scattered plot represents the weights of harvested tumors. Data are mean ± SEM, n=14; unpaired t-test; **p < 0.01. **(K)** H&E and IHC analysis with ECD and Ki-67. Representative images from three independent tumors (T1, T2 and T3) from both groups, in 400x magnification (scale bar, 200 µm).

To assess the impact of ECD OE on *in vivo* oncogenesis of PC cells, we introduced a mCherry-luciferase reporter into vector control and ECD-overexpressing LNCaP cells and subcutaneously injected these in the dorsal flank region of 8-weeks old male athymic nude mice. Tumor growth was monitored twice a week using calipers and weekly using bioluminescence imaging. Compared to the time course of tumor growth in mice implanted with control cells, those implanted with ECD-overexpressing cells developed more tumors that appeared earlier and grew to be significantly larger (**Fig. 5F-J, Table S2).** IHC staining of tumor sections confirmed the overexpression of ECD in ECD-OE tumors **(Fig. 5K)**. H&E-stained tumor sections showed that ECD-OE tumors had more necrosis **(Fig. 5K)**, a characteristic of aggressive tumors [41]. Finally, ECD-OE tumors showed increased Ki-67 staining **(Fig. 5K)**, a marker of tumor cell proliferation [42]. Together, the combination of *in vitro* and *in vivo* analyses demonstrates that ECD OE enhances the tumorigenic behavior of PC cells.

### ECD overexpression in prostate cancer cells upregulates the glycolysis pathway

To explore how ECD OE promotes the tumorigenic behavior of PC cells, we conducted RNA-seq analysis of xenograft tumors formed by ECD-overexpressing vs. Vector-expressing LNCaP cells followed by bioinformatics analyses. Principal component analysis (PCA) demonstrated the distinct clusters of the four each of the two types of tumors **(Fig. 6A**). Based on a log fold change >1 and false discovery rate (FDR) of <0.05, we found 520 differentially expressed genes (DEGs) in ECD-overexpressing compared to vector control tumors. Gene set enrichment analysis (GSEA) using the hallmark gene sets showed a significant upregulation of hypoxia, glycolysis, angiogenesis, myogenesis, apical junctions, TNFα signaling via NFkB and mTORC1 signaling in ECD-OE tumors compared to vector control tumors (**Fig. 6B & C**). These signaling pathways are well-known to be upregulated in aggressive prostate cancers [43–45]. Among the genes upregulated in ECD-OE LNCaP tumors were key glycolytic pathway genes (*ALDOA, LDHA, PKM, PGK1, HK1, HK2, ENO1*), Epithelial to Mesenchymal Transition (EMT) related genes (*VIM, SNAI1, TWIST2, TGFb1, TGFb2, HMGA2),* invasion and migration associated genes (*MMP2, MMP13, MMP14, MMP15* and *MMP24*), and genes associated with Notch signaling (NOTCH1, NOTCH2, NOTCH3 and NOTCH4), FGFR signaling (SPRY2, FGFR1) [44], and a previously identified aggressive prostate cancer signature gene set (*CCND2, RNF19A, DPP4* and *PRKDC*) [45] (**Fig. 6D**). In addition, ECD-OE tumors exhibited increased expression of neuro-endocrine prostate cancer (NEPC) related genes (*SYP*, *EZH2, ENO2, NRP2, NCAM1 and SOX2)*, a features of aggressive metastatic castration resistant PC (mCRPC) [46] (**Fig. 6D**). qRT-PCR analysis of mRNA isolated from xenograft tumors and *in vitro* cultured LNCaP cell lines validated the significant upregulation of glycolytic genes *HK2, LDHA and PKM2* (**Fig. 6E-G**) and NEPC related *SYP, NCAM1* and *ENO2* **(Fig. S7A & B)** in ECD-OE group compared to vector control. To further validate our qRT-PCR analysis of glycolytic mRNAs at protein level, we performed IHC analysis of vector and ECD-OE xenograft tumors and observed higher staining of HK2 and LDHA in tumor sections of ECD-OE LNCaP xenografts, as compared to controls (**Fig. S8)**.

**Fig. 6.**
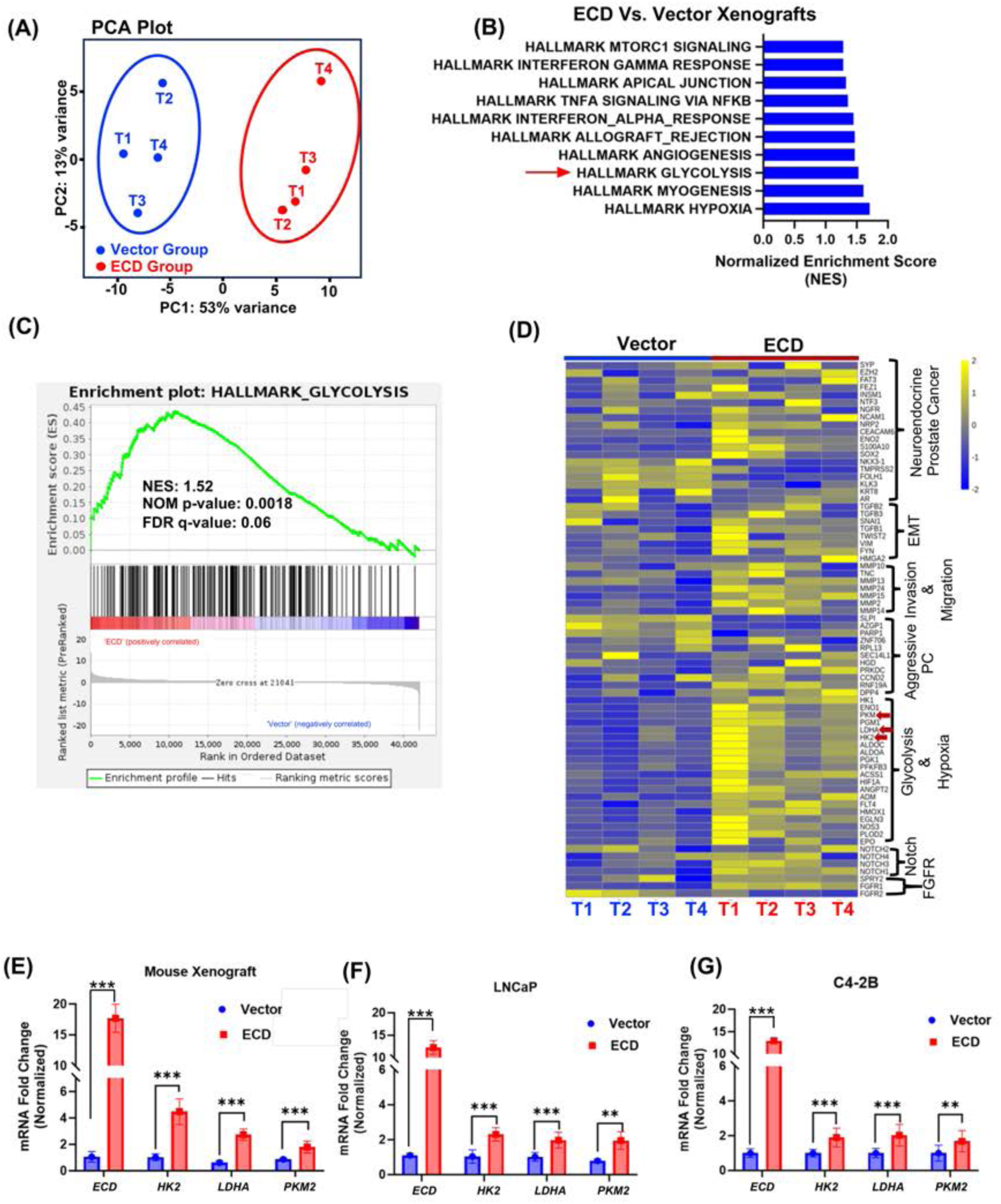
ECD regulates mRNA expression of glycolytic and neuro endocrine signature genes. **(A)** Principal component analysis (PCA) analysis of RNA-seq data from LNCaP-vector expressing and LNCaP-ECD-OE xenograft tumors shows distinct clustering of datasets. **(B)** Bar plot displays enriched hallmark GSEA (Gene set enrichment analysis) gene sets. X-axis represents normalized enrichment scores (NES) of the signaling pathways with significant nominal p-values (NOM p-val) and <25% false discover rate q-values (FDR q-val). Red arrow indicates glycolysis (followed further in this study). **(C)** Enrichment plot of glycolysis in ECD-OE xenograft tumors compared to vector expressing xenograft tumors. NES, NOM p-val and FDR q-val. are indicated on the plot. **(D)** Heatmap of key glycolytic genes and indicated signature genes of PC oncogenic pathways is presented. Yellow represents upregulated and blue as downregulated genes. Major glycolytic genes are marked as red arrows. **(E-G)** qRT-PCR confirms the overexpression of *ECD* in xenograft tumors and prostate cancer cell lines. 18s rRNA was used for normalization. qRT-PCR analyses of indicated glycolytic genes. mRNA quantitation data represents mean ± SEM with two-tailed unpaired *t* test. n = 3; **, p < 0.01; ***, p< 0.001.

Together, these results suggested the upregulation of glycolysis as a potential mechanism of ECD-promoted oncogenesis, consistent with the previously identified role of ECD in promoting glycolysis related gene expressions in mammary tumors of *ECD*Tg mice [23].

Given our’[18, 19, 47] and others’ recent findings that ECD regulates various aspects of mRNA biogenesis [48] and our recent findings that ECD is a RNA binding protein [47], we examined if ECD directly binds to and regulates glycolytic genes. LDHA and PKM2 were among the ECD interacting mRNAs identified through our recent analyses using enhanced crosslinking and immunoprecipitation (eCLIP) assay [47]. Using an RNA immunoprecipitation (RIP) assay, we found that *LDHA*, *PKM2* and *HK2* mRNAs were associated with ECD in both LNCaP **(Fig. 7A)** and C42-B **(Fig. 7B)** cell lines. Next, we analyzed the stability of these mRNAs in control vs. ECD-overexpressing cells by determining mRNA half-lives following treatment with actinomycin-D. These analyses revealed significantly longer (1.5 to 2-fold) half-lives of *PKM2*, *HK2* and *LDHA* in ECD-overexpressing vs. control LNCaP **(Fig. 7C-E)** and C4-2B **(Fig. 7F-H)** PC cells, determining that ECD binds to and stabilizes the mRNAs of key glycolytic genes.

**Fig. 7.**
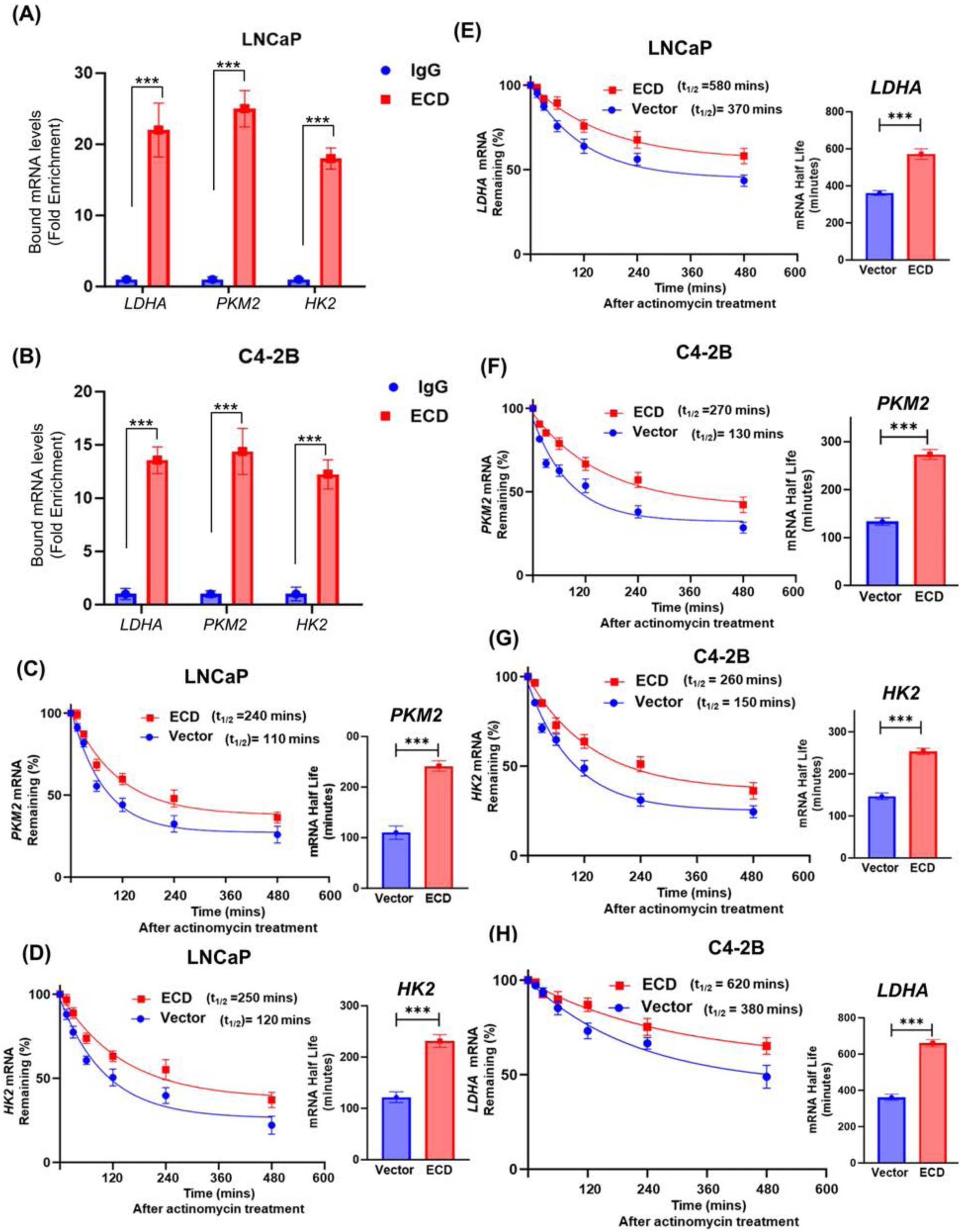
ECD is associated with mRNAs of glycolytic enzymes and regulates their stability. **(A & B)**. Indicated PC cells were subjected to RNA immunoprecipitation using IgG or anti-ECD antibodies and processed as per instructions in Magna RNA RIP kit. Co-precipitated RNAs were purified and the indicated target RNAs were detected by qRT-PCR using gene specific primers. Graph display mean fold enrichment in each respective gene, as compared to IgG control. **(C-H)** mRNA stabilities of various glycolytic enzymes after actinomycin D (5 µg/mL) treatment for indicated time points in vector or ECD expressing PC cells. Trend lines were created by measuring relative abundance of each transcript with respect to the 0-minute time point for vector and ECD-OE LNCaP and C4-2B cells and are expressed as percentage of mRNA remaining. Transcript abundance was measured by qRT-PCR using 18S rRNA as a housekeeping gene. Half-life (t_1/2_) for indicated mRNAs are displayed for vector and ECD-OE cells. Bar graphs display mean fold enrichment in each respective gene, as compared to IgG control. Quantitation data represents mean ± SEM with two-tailed unpaired *t* test. n = 3; ***, p< 0.001.

To functionally link the upregulation of glycolytic gene mRNA levels upon ECD OE to glycolysis, a Warburg effect [49], we measured glucose uptake and glycolytic rates. Indeed, there was a significant increase in glucose uptake in ECD-OE LNCaP (**Fig. 8A**) and C4-2B (**Fig. 8B**) cell lines as compared to vector expressing cells. To directly assess the impact on glycolysis, we conducted Seahorse XFe glycolytic rate assays to determine the proton efflux rate (PER) from glycolysis (GlycoPER) and extracellular acidification rate (ECAR) as readouts of glycolysis. Notably, the PER (**Fig. 8C & D**) and ECAR (**Fig. 8E & F**), measured in the presence of rotenone (Rot) and antimycin A (AA) to inhibit mitochondrial respiration, were increased in ECD-OE cells compared to vector cells. Additionally, ECD-OE cells showed increased levels of basal glycolysis (**Fig. 8G & H**) as well as of glycolytic reserve/compensatory glycolysis (**Fig. 8I & J**) after adding 2-Deoxy-D-glucose (2-DG) compared to their vector controls. Further, cellular oxygen consumption rate (OCR) was increased in ECD-OE PC cells relative to their controls **(Fig. S9A & B)**. Since increased glycolysis is often associated with enhanced glycogen synthesis [6], we performed Periodic Acid–Schiff (PAS) staining to assess glycogen deposition, [50] along with IHC staining for glycogen synthase 1 (GYS1) and glycogen phosphorylase (PYGB), key enzymes involved in glycogen metabolism,[50] in tumor sections from Vector and ECD-OE LNCaP xenografts. Notably, we observed OE of ECD increased glycogen synthesis and metabolism, as demonstrated by higher staining intensities of PAS **(Fig. S10A)**, GYS1 and PYGB **(Fig. S10B)**. These results demonstrate that ECD OE in PC cells increases the expression of key glycolytic genes through binding to and stabilization of their mRNAs to promote the Warburg effect.

**Fig. 8.**
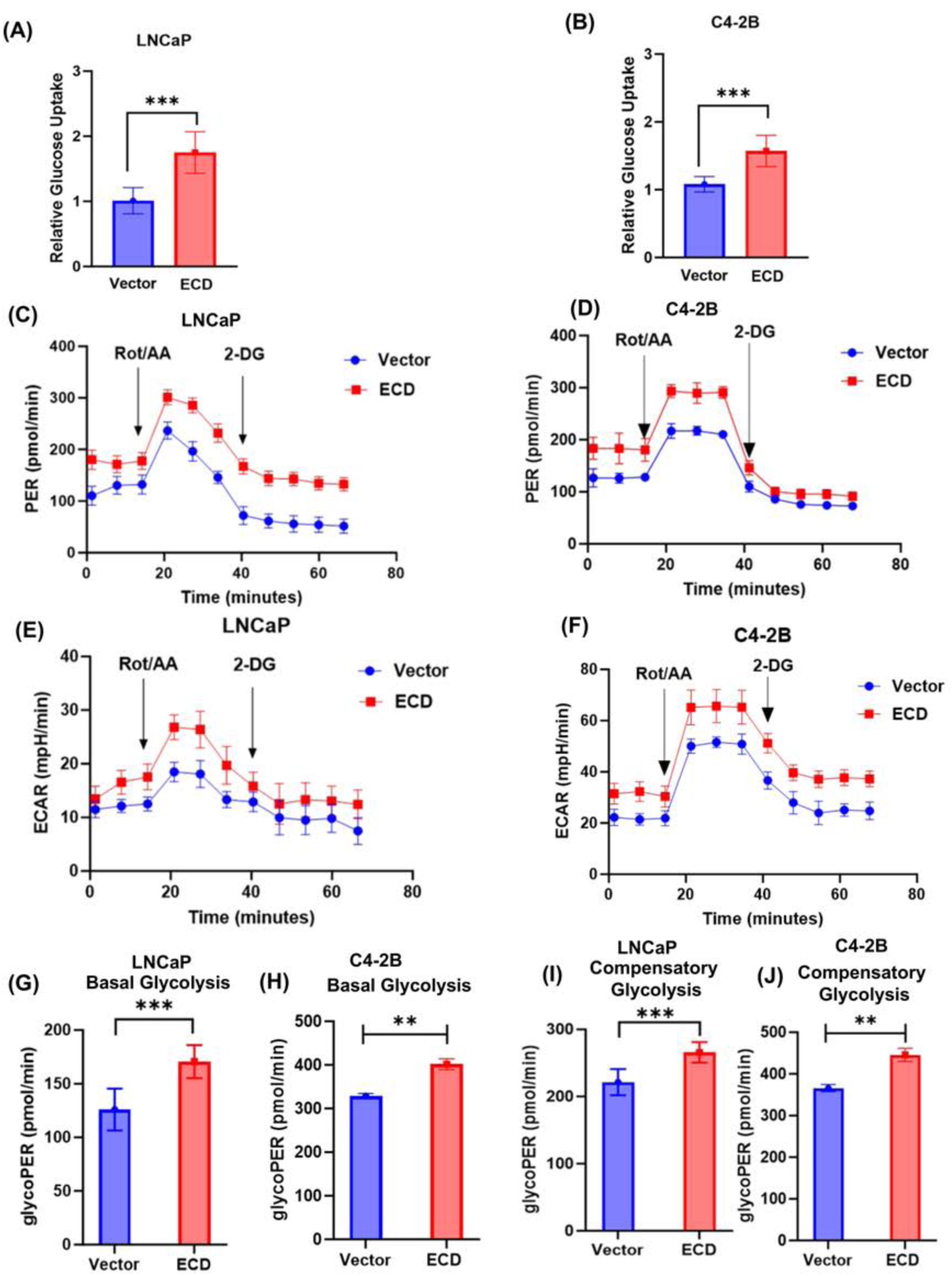
ECD overexpression increases glucose uptake and glycolytic rate. Glucose uptake in LNCaP **(A)** and C4-2B **(B)** PC cell lines. The values were normalized with respective to cell counts and depicted as compared with vector control. Quantification of results from three biological experiments, each with six technical replicates is shown as a bar graph. Data represents mean ± SEM and two-tailed unpaired Student’s t-test (*p < 0.05, **p < 0.01, ***p < 0.001). **(C-J)** Sea horse glycolytic rate was assessed in vector and ECD-OE LNCaP **(C)** and C4-2B **(E)** cells. Cells were seeded in 96-well plates and exposed to Rot/AA (rotenone and antimycin cocktail) and 2-DG to measure proton efflux rate **(PER, C & D)** and extracellular acidification rate **(ECAR, E & F)**. One representative PER and ECAR plots from three independent experiments are displayed (**C-F**). Basal glycolysis **(G & H)** and compensatory glycolysis **(I & J)** rates are presented as bar graphs, glycoPER in pmol/min calculated from subtracting ECAR (extra cellular acidification rate) from PER (proton efflux rates) (**G-J**). Quantitation data represents mean ± SEM with two-tailed unpaired *t* test. *n* = 3; **, p < 0.01; ***, p < 0.001.

## Discussion

While androgen signaling through AR is universally accepted as an early driver of prostate tumorigenesis with well-established targeting agents, androgen and AR directed therapies eventually fail in most patients [51]. Metabolic reprogramming represents a key molecular determinant of the aggressive behavior of PC and therapeutic resistance, with enhanced glucose utilization through aerobic glycolysis (Warburg effect) as a major factor [49]. Molecular pathways that link AR signaling and glycolytic metabolism in PC are of interest as these are potential biomarkers, as well as for therapeutic targeting in conjunction with AR directed therapies. Here, we identify the Ecdysoneless (ECD) protein as a novel link between AR signaling and promotion of glycolysis and demonstrate that ECD, which is overexpressed in >40% of PC patient tumors and predicts shorter survival, is an AR-responsive target gene whose increased expression promotes glycolysis through ECD binding to glycolytic gene mRNAs to enhance their stability. Our studies elucidate a novel mechanism to promote metabolic reprogramming and tumorigenesis in PC and raise the possibility of glycolysis pathway co-targeting with AR in ECD-OE PC.

AR-mediated signaling is a critical element of the early progression of all PCs and remains an oncogenic driver in most late-stage PCs [48]. Our comprehensive analysis establishes that ECD OE is directly regulated by AR signaling. We found a strong positive correlation between AR and ECD mRNA expression levels across PCs (TCGA-PRAD dataset) **(Fig. 1A)** and cell-based analyses of PC cell lines showed a dose- and time-dependent upregulation of ECD mRNA and protein levels by exogeneous androgen treatment of PC cells **(Fig. 1B-I)** with siRNA KD demonstrating a requirement of AR for basal as well as androgen-stimulated ECD expression **Fig. 1J & K**),. Direct evidence for ECD as a novel androgen-responsive AR target gene in PC was provided by ChIP **(Fig. 2A-D)** that established the potential AREs identified by in-silico analysis of the ECD promoter to be authentic AR binding sites and mutational analyses of the three AREs using a luciferase reporter assay **(Fig. 2E & F)** demonstrated their combined requirement for androgen-induced ECD expression. Consistently, treatment of PC cell lines with anti-androgen Enzalutamide decreased ECD levels **(Fig. 2H-K)**. Further the role of ECD in PC was established by genetic loss as well as gain of function approaches in PC cell line models. Inducible ECD KO led to a marked reduction in key *in vitro* oncogenic traits **(Fig. 3A-M)**.

While ECD plays a physiological role in cell cycle progression and ER stress mitigation, and its overexpression correlates with poor prognosis and shorter survival in breast, pancreatic and gastric cancers [21, 22], any role and regulation of ECD in PC has remained unexplored. Our IHC analyses of a large cohort of PC patient tumors using a well-authenticated anti-ECD monoclonal antibody [52], showed ECD OE in nearly 43% of PC cases and such overexpression specified shorter patient survival (**Fig. 4A-E**). Analysis of publicly available TCGA PC adenocarcinoma dataset demonstrating shorter survival of PC patients with higher ECD mRNA expression **(Fig. 4F)** lend further support to our IHC results. Notably, high ECD OE was associated with key bad prognostic indicators like higher Gleason score, advanced tumor stage and positive resection margins **(Table S1)**. Thus, ECD OE emerges as a novel biomarker of poor prognosis and shorter patient survival in PC.

These results support a novel mechanism whereby AR signaling induces the upregulation of ECD expression in PC as one pro-oncogenic pathway. It is noteworthy that while we and others have reported the upregulation of ECD expression in other cancers, the mechanisms responsible for such overexpression remain unknown. Given the androgen/AR mediated upregulation of ECD expression in PC identified here, it will be of great interest to assess the potential regulation of ECD OE across cancers through other nuclear hormone receptors, especially in hormonally regulated malignancies. Notably, the AR requirement for ECD OE was observed in an androgen-dependent as well as an androgen-independent PC cell line model, suggesting that AR signaling rather than exogenous androgen requirement is a key determinant of ECD OE in PC as significant subsets of CRPCs are AR-independent [49]. Intriguingly, elevated levels of ECD are observed in the AR-negative PC cell line PC3 (**Fig. S1A**), indicating that ECD regulation may not be solely dependent on AR signaling. The possibility remains that ECD expressions in AR-negative PC may be regulated by alternate steroid hormone receptor signaling pathways, such as those mediated by glucocorticoid receptors (GR). Literature suggests that hormone responsive elements (HREs) are similar to androgen response element (ARE) [53], and AR-negative PC often exhibit higher GR expression [54].. Future studies utilizing both AR-positive and AR-negative PC will help determine the role of ECD through GR-mediated signaling.

Besides emerging as a correlative biomarker of aggressive PC, our studies clearly demonstrate that ECD OE is biologically important in promoting the tumorigenic behavior of PC cells **(Fig. 5A-E, and Fig. S6).** Notably, both vector and ECD-OE PC cells remain hormone responsive for their proliferation **(Fig. S6N)**. Importantly, ECD OE imparted a more aggressive *in vivo* tumorigenic phenotype with earlier growth of xenografted tumors, significantly larger tumors (**Fig 5F-J**), and higher levels of Ki-67 staining and necrotic lesions indicative of hyper-proliferative and rapidly growing aggressive tumors (**Fig. 5K**), consistent with known association of hyperproliferation and necrosis with aggressive prostate tumors in patients [55]. Thus, like the role of ECD OE in other cancers such as breast cancer [23], its overexpression appears to propel PC towards increased aggression. We note that while our studies in established PC cell lines clearly reveal the impact of ECD OE to promote tumorigenic phenotype, our studies do not address any potential role of ECD OE in early malignant transformation driven by androgen-AR axis. In our previous studies using immortalized mammary epithelial cell models, ECD OE alone did not impart a tumorigenic phenotype [56]. However, since ECD plus mutant RAS co-overexpression in immortal mammary epithelial cells promoted full tumorigenesis [56], and transgenic overexpression of ECD in the mammary epithelium led to hyperplasia and late-onset tumor induction [23], incorporation of ECD transgene or tissue-specific ECD KO genetic models of prostate tumorigenesis should help assess a potential early pro-tumorigenic role of ECD in PC.

Our mechanistic studies provide a plausible explanation for the pro-oncogenic role of ECD OE in PC. Our RNA-seq analyses identified a potential role of ECD OE in upregulating the mRNA levels of key glycolytic genes, *HK2, LDHA* and *PKM2* **(Fig. 6)**. Consistent with our recently established biochemical function of ECD as an RNA-binding protein [47], these mRNAs are physically associated with ECD **(Fig. 7A & B)** and ECD OE enhances their stability **(Fig. 7C-H).** A significant increase in glucose uptake (**Fig. 8A & B**) and glycolysis measured with Seahorse glycolytic assays (**Fig. 8C-J**) directly established that the increased expression of glycolytic gene mRNAs upon ECD OE translates into function of ECD in glycolysis. Furthermore, IHC staining of ECD-OE LNCaP tumor xenografts showed higher expression of LDHA and HK2 (**Fig. S8**), along with increased Periodic Acid Schiff (PAS) staining, and an increase level of GYS1 and PYGB (proteins involved in glycogen metabolism) (**Fig. S10A & B**) further underscoring the role of ECD to regulate glucose and glycogen metabolism. Thus, we posit that increased expression of ECD in PC, initiated by androgen-AR signaling, serves as one mechanism to promote a glycolytic switch to enhance the aggressiveness of PC. This mechanism is consistent with our analyses of ECD transgene driven mammary tumors, which also exhibited an upregulation of glycolysis enzyme mRNAs and increased glucose uptake [23]. A linkage of ECD OE to increased glycolysis as a mechanism to enhance PC tumor aggressiveness is also consistent with prior studies that have demonstrated a key role of aerobic glycolysis in promoting cell survival [57], metastasis [58] and drug resistance in PC [59]. While our studies suggest the ECD OE as a potential positive regulator of glycolysis in >40% of prostate tumors, clearly other mechanisms are likely to play a role either concurrently with ECD OE or in patients without ECD OE. Notably, AR signaling is known to control glycolysis directly in PC, [60] however other mechanisms like aberrant FGFR tyrosine kinase signaling [61] and activation of Wnt/β-catenin signaling [62] have also been linked to enhanced glycolysis in PC. Future studies will be needed to assess how ECD OE may synergize with other mechanisms that promote glycolysis. Additional investigations are needed to elucidate whether ECD, as an RNA-binding protein, and regulator of RNA splicing contributes to splicing modulation of glycolytic genes, to further enhance glycolytic activity.

In conclusion our work reveals ECD as a novel androgen-regulated AR target gene and demonstrates that ECD OE, a feature of >40% PC patients, promotes PC tumor aggressiveness through upregulation of glycolysis. Our findings open the possibility of exploring glycolysis-targeted therapeutic approaches for PC patients with high ECD expression. These results support oncogenic role of ECD in both hormone dependent and hormone independent AR-expressing prostate cancer.

## Materials and Methods

### Antibodies and reagents

Antibodies: anti-AR (Invitrogen cat # MA5-13426), β-actin (Sigma cat # A5441), Ki67 (Abcam cat # ab16667), anti-PYGB (Proteintech cat # 12075-1-AP), anti-LDHA (R&D cat # MAB9158), anti-HK2 (Abcam cat #ab209847), anti-GYS1 (ThermoFisher cat # 10566-1-AP), anti-ECD antibody generated by us has been described previously [13]. 5α-Dihydrotestosterone solution (Millipore Sigma cat # D-073), Chemiluminescence signals were detected by (ECL Plus; Amersham-Pharmacia cat # NEL-105001EA) and Matrigel (Coring Cat # 356230), and Isoflurane (cat # 502017) from MWI Animal Health.

### Cell lines and medium

PC cell lines LNCaP, C4-2B, 22Rv1, VCaP, PC-3 and HPV18 immortalized prostate epithelial cells (RWPE-1) were obtained from the American Type Culture Collection (ATCC). Cancer cell lines were maintained in complete RPMI medium with 10% fetal bovine serum [63] and RWPE-1 was cultured in keratinocyte growth medium (KGM Bullet Kit, Lonza) [19]. For androgen deprivation experiments, cells were cultured in phenol red-free complete RPMI medium supplemented with 10% charcoal stripped FBS (cat # S11650 Atlanta Biologicals). Lenti-X 293T cell line used for lentiviral packaging was from Takara Bio (cat # 632180) and cultured in complete DMEM medium [63]. Cells were cultured in a humidified atmosphere with 5% CO2 at 37°C. Cell lines were regularly evaluated for mycoplasma and continuously cultured for no more than 3 months.

### Chromatin Immunoprecipitation assay

Cells grown in charcoal stripped medium and were treated with DHT for 3 h, washed with PBS and treated with 1% formaldehyde for 10 min to crosslink protein-DNA complexes. Cells were rinsed and processed for ChIP with control IgG (Abcam, cat # ab172730) or anti-AR antibody (Abcam, cat # ab185913). The following PCR Primers were used to determine the enrichment: ECD AR Binding site Forward primer -TTAATTCTGGGATCTTTATCTGTAG; ECD AR Binding site Reverse primer-GGCTCCAGGAAAGTGCTCACTTGAC. Real-time PCR quantification was performed in triplicates using SYBR Green PCR master mix (Applied Biosystems cat # 4309155). AR Binding percentage was calculated post normalization against IgG.

### Luciferase-promoter reporter assays

The promoter sequences of GAPDH (Genome=hg38;chr12+: 6533244-6534713; TSS=6534517; Upstream=1273, Downstream=196; Length=1470), and wildtype (WT) ECD (Genome=hg38;chr10-:73169381-73167884; TSS=73168095; Upstream=1286, Downstream=211; Length=1498) or its predicted AR binding site deletion mutants (Details in **Table S3**) were synthesized and cloned upstream of the Gaussia luciferase in the lentiviral pEZX-LvPG04 vector (sequence-confirmed constructs were custom-made by Genecopoeia; cat# HPRM35342-LvPG04-WT, GAPDH-LvPGO4). The ECD promoter mutants and the sequences deleted from TSS are listed in **Table S3**. The reporter constructs were introduced into LNCaP and C4-2B cell lines using lentiviral supernatants. For promotor luciferase experiments, transduced cells were cultured in charcoal stripped serum containing medium for 5 days and then treated with 100 nM DHT for 3 h. Post treatment, supernatant was collected and analyzed using the Secrete-Pair Dual Luminescence Assay Kit (cat # LF031) to measure luciferase activity following the manufacturer’s protocol. The fold induction was calculated after normalizing the values against secreted alkaline phosphatase luminescence, used as an internal control.

### RNA isolation and quantitative real-time PCR

Total RNA isolation and qRT-PCR analysis was conducted in indicated PC cell lines as described previously [18]. The qRT-PCR primers are listed in Table **S4**.

### Generation of CRISPR-Cas9 mediated inducible ECD Knockout (KO) cell lines

To generate inducible ECD KO cell lines, Lentiviral supernatants were generated as mentioned above for Edit-R inducible Lentiviral hEF1a-Blast-Cas9 plasmid (cat # CAS11229) and Edit-R lentiviral plasmids carrying guide RNAs against non-targeting control (NTC) (cat # GSG11811), or human ECD gRNA1 (cat # GSGH11838-246505144, Seq-TCCAAAGTCTCAACCCACAA) or gRNA2 (cat # GSGH11838-246505139, Seq-CTTGGGTATACTTACCCTAT). All plasmids were procured from horizon discovery. Cells were transduced first for cas-9 expression, then selected in medium containing 5 μg/ml blasticidin for 5 days. Inducible Cas-9 expressing cells were then transduced with either NTC or gRNA1/2 lentiviral supernatants and selected in 5 μg/ml blasticidin + 1 μg/ml puromycin for additional 5 days.

### Transwell migration and invasion assays

Cell lines were subjected to serum deprivation (cultured in complete RPMI medium with 0.5 % FBS) for 24 h, trypsinized and seeded in transwell migration (Corning BioCoat 8.0 μm cat # 354578) or invasion (BioCoat Matrigel 8.0 μm cat # 354480) chambers at 100,000 cells per well in 0.5% FBS containing medium. After three hours of incubation at 37°C, complete RPMI medium containing 10% FBS was added as a chemoattractant in the bottom chambers. After 24 h of incubation, the cells on the top of filters were scrapped off and the migrated cells on the bottom of filters were fixed in ice-cold methanol, stained with crystal violet, and rinsed in PBS. Cells in six random fields per well were imaged at 100x magnification and quantified using the ImageJ software. Experiments were repeated three times independently with triplicates each time.

### Anchorage-independent growth assays

20,000 vector-control or ECD-OE cells in complete RPMI medium containing 0.3% agarose were seeded on top of a 0.6% bottom layer of agarose in 6-well plates, as described [56]. Cultures were fed every 2 days. Twenty-one days after cell seeding, the colonies were stained with 0.05% crystal violet in 25% methanol. Colonies were imaged under 10x magnification with phase contrast microscope and counted using the image J software. Experiments were performed in triplicates, and each experiment was repeated three times.

### Matrigel organoid cultures

2,000 vector-control or ECD-OE cells resuspended in the medium containing 4% Matrigel were seeded per well on top of a base layer of 100% Matrigel in eight-well Nunc chamber slides. The cells were allowed to form organoid colonies for 6 days and then imaged. The ImageJ software was used for quantification with a size cut-off of 70 μm in diameter. All experiments were performed in triplicate wells and repeated three times.

### Prostate cancer patient tissue microarrays and immunohistochemical analysis

The prostate cancer tissue specimens were obtained with informed consent from patients who underwent radical prostatectomy at the Department of Urology, Institute for Pathology, University Hospital “Carl-Gustav-Carus”, University of Technology, Dresden between 1999 and 2005. Patient age ranged between 43 and 74 years (median 60 years). Preoperative PSA levels ranged from 0.8 to 39 ng/ml (median 7.2). Clinical follow-up data was assessed annually. The distribution of Gleason scores (GS) in the cohort was as follows: GS 2–6: 192 (34.5%), GS 7-10: 364 (65.5%). Three hundred and eighty-six cases had organ-confined carcinomas (pT2); 170 cases showed extracapsular tumor extension (pT3). The surgical margins were clear (R0) in 396 cases; 157 cases had positive margins (R1). The median follow-up time of all cases was 60 months. Tissue microarrays (TMAs) composed of 556 cases of paired normal/hyperplastic prostate and prostate cancer (one core per case) were analyzed for ECD expression using an anti-ECD monoclonal antibody [13]. 542 samples were classified as normal, 401 samples were hyperplastic, and 258 samples were classified as high-grade PIN. We have followed data of 522 prostate cancer patients (each patient was represented only once, so 556 tumors are from 566 different patients). The intensity of staining was expressed as a semiquantitative scoring system: negative = 0, low = 1, moderate = 2 and High =3. Stratification into ECD-High vs ECD-Low expression status reflects cases with staining intensity of 0 or 1 vs. 2 or 3, respectively. Kaplan-Meier survival analysis compared 224 ECD-High (2-3) vs. 298 ECD-Low (< or = 1) cases. A pathologist evaluated all slides. ECD expression (low vs. high), tumor grade (Gleason 1–6 vs. 7–10), pT-stage (pT1/2vs. pT3/4), R-status (R0 vs. R1), age (<60 years vs. >60 years) and pre-operative PSA (low/<7.2 vs. high/>7.2 ng/ml) were used in multivariate Cox regression analysis. The Charite approved the use of PC tissue samples by the University Ethics Committee under the protocol ‘Retrospective analysis of tissue samples by immunohistochemistry and molecular biological techniques’ (EA1/06/2004) on 20 September 2004.

### ECD overexpression in prostate cancer cell lines

The ECD (GeneID: 11319) CDS in the mammalian expression vector pLenti-C-Myc-DDK-P2A-Puro and empty Vector control plasmids were obtained from (Origene Biotech cat # PS100092). Lentiviral supernatants were generated by transient co-transfection of Lenti-X 293T cells with ECD expression vector and packaging plasmids psPAX2 (Addgene, cat # 12260) and pMD2.G (Addgene cat # 12259) using X-treme GENE HP DNA transfection reagent (Roche cat # 06366236001). Collected lentiviral supernatant was added to PC cells in the presence of polybrene (8 µg/ml, Sigma # H9268). Post-infection, transduced cells were selected in 1 μg/ml puromycin for 5 days, and ECD levels assessed by Western blotting using an anti-ECD monoclonal antibody [13]. For siRNA KD, cells were transfected with 40 nM of control or gene-specific siRNAs (AR siRNA, Santa Cruz, cat # SC29204) using the DharmaFECT 1 transfection reagent (cat # T-2001-03, Dharmacon, Pittsburgh, PA).

### Cell proliferation assay

2,000 cells were plated per well in 96 well plates, with six replicates for each condition. Medium was changed on alternate days. Cell proliferation was measured at the indicated times using the Cell Titer-Glo® luminescent cell viability assay (Promega, cat # G7570) following the manufacturer’s protocol.

### Xenograft studies and IVIS imaging

All animal experiments were pre-approved by the UNMC Institutional Animal Care and Use Committee (IACUC Protocol # 22-059-10-FC). To assess the effect of ECD OE on tumorigenesis, 3×10^6^ vector control or ECD-OE LNCaP cells engineered with lentiviral mCherry-luciferase in 0.1 ml 50% Matrigel (BD Biosciences) in cold PBS were injected subcutaneously in 8-week-old athymic nude male mice (n=14) on both sides of dorsal flank. Tumor growth was monitored twice a week using calipers and once a week by IVIS bioluminescence imaging for up to 37 days. Tumor volume was calculated as length x width x depth/2 [23]. Mice were euthanized when mice reached pre-determined humane endpoints or reached 2 cm^3^ in volume, as per the institutional IACUC guidelines. For IVIS imaging, mice received 200 μl D-luciferin (15 mg/ml in PBS; Millipore Sigma #L9504) intraperitoneally 15 min before isoflurane anesthesia and were placed dorsoventrally in the IVIS™ Imaging System (IVIS 2000). Images were acquired using the IVIS Spectrum CT and analyzed using the Living Image 4.4 software (PerkinElmer). The fold change in total Photon flux (p/s) over time was used to compare tumor growth between the vector and ECD-OE cells groups.

### Immunohistochemistry of mice tumors

The mouse tumors harvested at necropsy were fixed in 10% neutral buffered formalin (cat # 316-155; PROTOCOL), paraffin-embedded and 5 mm sections deparaffinized in xylene and rehydrated in descending alcohols followed by IHC staining of indicated antibodies as previously described [23]. Periodic Acid Schiff’s staining of tumor sections was conducted in tissue science core facility of the University of Nebraska Medical Center.

### RNA sequencing, bioinformatics, and statistical analysis

Trizol-isolated total RNA from the core piece of the isolated tumors was analyzed on a Bioanalyzer for quality and purity. RNA-seq libraries were prepared and subjected to 100 bp paired-end sequencing on an Illumina NovaSeq 6000 system (UNMC Genomics Core). Analysis of RNA-seq results was done using the nf-core RNA-seq pipeline version 3.9 [64], accessed via Nextflow version 22.10.3 [65] and executed with Singularity [66] to ensure reproducibility and portability. Paired-end sequencing reads were aligned to the human reference genome (GRCh38) using the Spliced Transcripts Alignment to a Reference (STAR) software version 2.7.11a [67], incorporating Ensemble gene annotations for accurate mapping. Quantification of transcript abundance was performed using StringTie2 version 1.3.6 [68], with transcripts per million (TPM) normalization to facilitate comparison across samples. Differential expression analysis between vector and ECD-OE xenograft tumors was conducted using DESeq2 [69]. Differentially expressed genes were identified using criteria of an adjusted p-value < 0.05 and a minimum twofold change in expression. For functional interpretation of the differential expression results, we utilized the Gene Set Enrichment Analysis (GSEA) software from the Broad Institute [70]. Enrichment plots were generated to visualize the distribution of gene sets and to identify pathways significantly associated with the observed transcriptional changes.

### RNA Immunoprecipitation (RIP) assay

RIP assays were performed using the EZ-Magna RIP Kit (Millipore, cat #17-701). Briefly, cells were harvested and lysed using RIP lysis buffer with 1 freeze-thaw cycle. Cellular debris was removed by centrifuging the cell lysates at 13K for 15 min. 1 mg aliquots of clarified lysate (BCA method) were incubated overnight at 4°C with protein A Dynabeads bound to 2.5 μg of polyclonal rabbit anti-ECD (Proteintech, cat # 10192-1-AP) or control IgG (Abcam cat # ab172730) antibodies. Post incubation samples were subjected to five washes in the wash buffer. The bound RNAs were retrieved and subjected to qRT-PCR analysis using gene specific primers. 10% of lysates served as input controls.

### Analysis of mRNA stability

cell lines were seeded in 6-well plates (200,000 cells per well). 24 h after plating, 5 μg/ml actinomycin D (catalog # A1410; Sigma, St. Louis, MO) was added and total RNA isolated at the indicated time points using the TRIzol method. cDNA was synthesized using iScript™ gDNA Clear cDNA Synthesis Kit (Bio-Rad), as above, followed by qRT-PCR with the indicated gene specific primers.

### Glucose Uptake-Glo assay

Cells were seeded at 25,000 cells per well in 96-well plates. 24 h later, the cells were washed twice with PBS and cultured in glucose-free DMEM media (Thermo Scientific cat # 11966025) for 1 h (glucose starvation). Glucose uptake assay was then performed using the Promega Glucose Uptake-Glo (J1341) assay kit according to the manufacturer’s protocol. Parallel CellTiter-Glo assays were used for normalization of cell numbers.

### Seahorse XF Glycolytic Rate Assay

The Oxygen Consumption Rate (OCR), Extracellular Acidification Rate (ECAR) and Proton Efflux rate (PER) were measured using the Seahorse XFe Glycolytic rate assay Kit (Agilent Technologies; cat #103344-100) according to the manufacturer’s protocol. Briefly, 3.5 × 10^4^ cells were seeded per well in 96-well Seahorse XFe cell culture plates and incubated overnight. Cells were washed and incubated with XF assay medium supplemented with 1 mM pyruvate (Sigma; cat# S8636), 2 mM L-glutamine (Invitrogen; cat#25030081) and 10 mM glucose (Gibco; cat#A24940-01) for 1 h at 37°C in a CO_2_-free incubator. The OCR, ECAR and PER estimations were performed as per the manufacturer’s instructions. Basal OCR and ECAR were assessed in response to rotenone/antimycin A (0.5 μM) and 2-deoxy-D-glucose (2-DG; 50 mM). ATP rate was estimated in response to oligomycin (1.5 μM) and rotenone/antimycin A (0.5 μM). Basal glycolysis was calculated from non-glycolytic acidification (background) ECAR/PER. Glycolytic reserve (Compensatory) was calculated as the difference between glycolytic capacity and basal glycolysis.

### Statistical analysis

Statistical analyses were done using the GraphPad Prism 11 software. Significance between two groups was determined by the two-tailed Student’s t test. Comparisons among three or more groups were made using either one-way ANOVA or two-way ANOVA with post hoc multiple comparisons test. A p-value of ≤0.05 was considered statistically significant. Survival curves were analyzed using the Kaplan-Meier analysis with the log rank test. The results are expressed as the mean ± SEM from three independent experiments, done in triplicates. The type of statistical test used in individual experiments is indicated in figure legends.

## Acknowledgements

This research was funded by Pilot grants from the Fred & Pamela Buffett Cancer Center (HB & VB); Department of Defense grants W81XWH-17-1-0616 and W81XWH-20-1-0058 to HB and W81XWH-20-1-0546, HT94252410337; and NIH grants R21CA241055 and R03CA253193 to VB; the NIH Pathway to Independence Award R00 GM1287671and the NIH MIRA award R35 GM147467 (to M.J.R.); the Raphael Bonita Memorial Fund, and support to UNMC core facilities from the NCI Cancer Center Support Grant (P30CA036727) awarded to Fred & Pamela Buffett Cancer Center and from the Nebraska Research Initiative.

## Author Contributions

Conceptualization and design: M.R., A.R.R., B.B.K., T.E.R., F.O., S.M., M.J.R., G.K., K.D., B.C.M., H.B., and V.B.

manuscript writing; M.R., A.R.R., B.C.M., T.E.R., K.D., M.J.R., H.B., and V.B.

data analysis and interpretation. M.R., A.R.R., B.B.K., T.E.R., F.O., S.M., M.J.R., G.K., K.D., B.C.M., H.B., and V.B.

manuscript editing; M.J.R., B.C.M., H.B., and V.B.

bioinformatics analysis for the manuscript. M.R., T.E.R., M.J.R.,

All authors have read and approved the final version of the manuscript and agree with the order of presentation of the authors.

## Data Availability Statement

The RNA-seq files generated in this study are available from the NCBI Gene Expression Omnibus (GEO) under the accession number: GSE287898.

---

**Table S1.** Multivariate Cox’s regression analysis of prostate cancer patient tumors with inclusion of cytoplasmic ECD expression. **RR: Relative risk; CI: Confidence interval.**

|  |  | Overall Survival |  |  |
| --- | --- | --- | --- | --- |
|  | No. of Cases | RR | 95%CI | p-value |
| **Cytoplasmic ECD Expression** |  |  |  |  |
| negative/weak | 315 | 1 |  |  |
| moderate/strong | 241 | 1.689 | 1.085-2.629 | 0.02 |
| **Tumour stage** |  |  |  |  |
| postoperative pT2 | 386 | 1 |  |  |
| postoperative pT3 | 170 | 3.126 | 1.899-5.146 | 0 |
| **Gleason** |  |  |  |  |
| Score 2-6 | 192 | 1 |  |  |
| Score 7-10 | 364 | 2.415 | 1.247-4.675 | 0.009 |
| **R-status** |  |  |  |  |
| R0 | 396 | 1 |  |  |
| R1 | 157 | 1.666 | 1.038-2.675 | 0.035 |
| **Age** |  |  |  |  |
| ≤60 Years | 194 | 1 |  |  |
| >60 Years | 362 | 0.971 | 0.934-1.011 | 0.151 |
| **PreOP PSA** |  |  |  |  |
| ≤7.2ng/ml | 279 | 1 |  |  |
| >7.2ng/ml | 269 | 1.529 | 0.965-2.425 | 0.071 |

**Table S2.** Incidence of tumor formation in vector or ECD-overexpressing LNCaP xenograft model.

| **No. of Days** | **LNCaP Control group** | **ECD OE LNCaP Group** |
| --- | --- | --- |
| 16 | 0 (0/26)/ **0%** | 6 (6/28)/ **21.4%** |
| 22 | 8 (8/26)/ **30.7%** | 20 (20/28)/ **71.4%** |
| 27 | 9 (9/26)/ **34.6%** | 23 (23/28)/ **82.1%** |
| 35 | 13 (13/26)/ **50%** | 26 (26/28)/ **92.8%** |

**Table S3.** List of ECD promoter mutants and the corresponding sequences deleted from TSS.

| **ECD Promotor Mutants** | **Sequence deleted from TSS** |
| --- | --- |
| CS-HPRM35342-LvPG04-01- site-1Δ | 698 bp – 682 bp |
| CS-HPRM35342-LvPG04-02-site-2Δ | 657 bp – 641bp |
| CS-HPRM35342-LvPG04-03-site-3Δ | 640 bp – 624 bp |
| CS- HPRM35342-LvPG04-04-site-1,2Δ | 698 bp – 682 bp, 657 bp – 641bp |
| CS-HPRM35342-LvPG04-05-site-1,3Δ | 698 bp – 682 bp, 640 bp – 624 bp |
| CS-HPRM35342- LvPG04-06 - site-2,3Δ | 657 bp – 641bp, 640 bp – 624 bp |
| CS-HPRM35342-LvPG04-07 - site-1,2,3Δ | 698 bp – 682 bp, 657 bp – 641bp,<br>640 bp – 624bp |

**Table S4.**
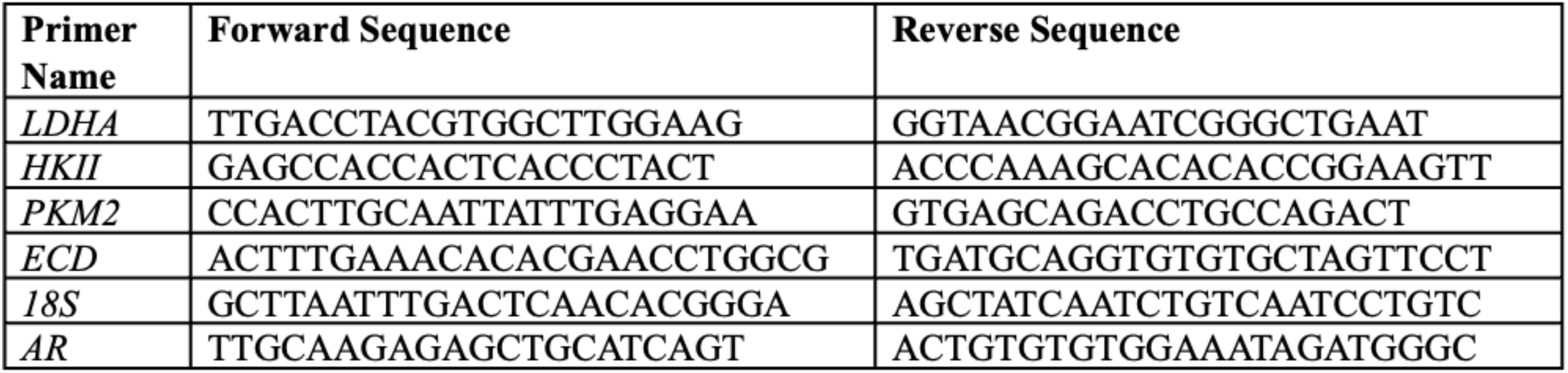
List of real time qRT-PCR primers of human genes.

**Fig. S1.**
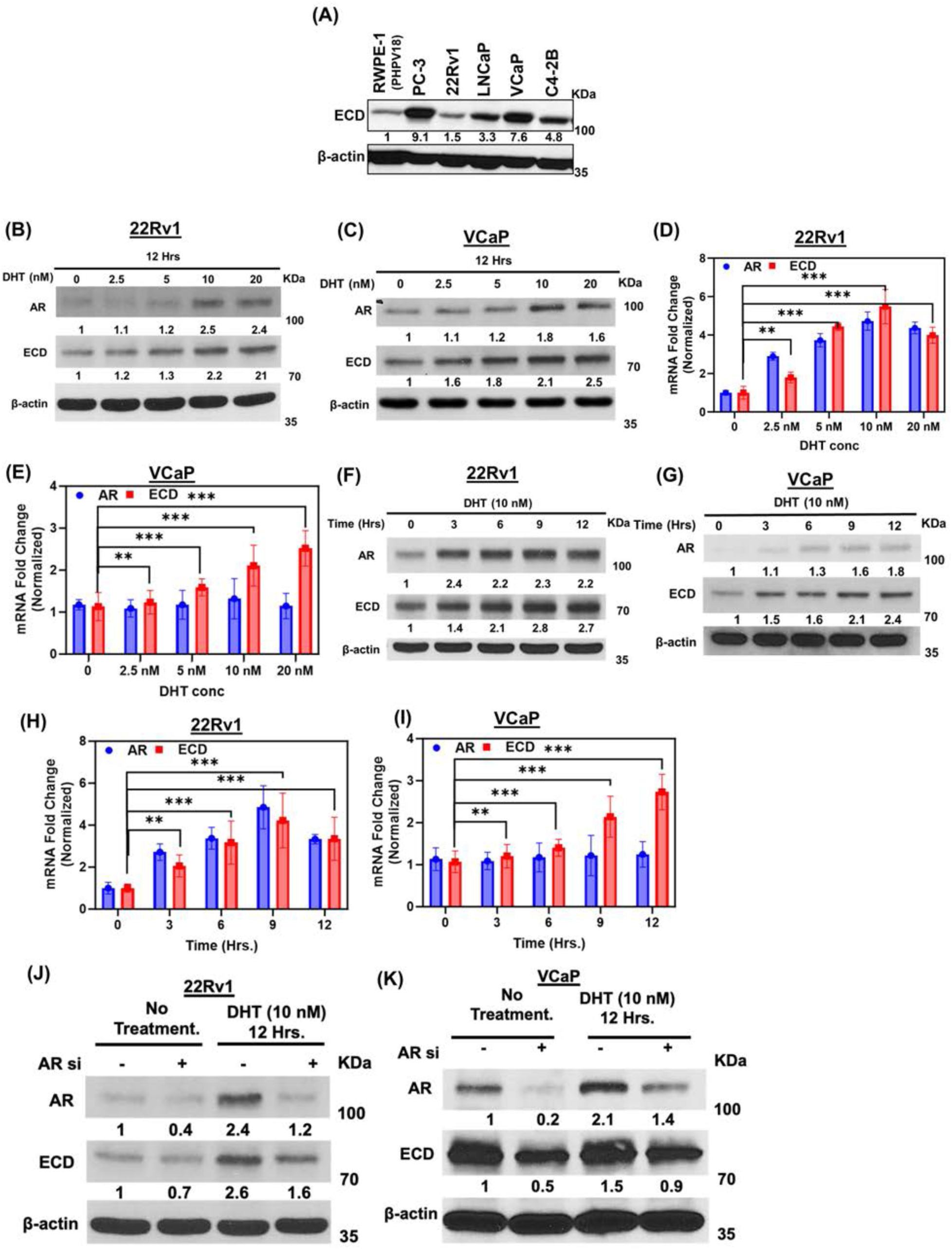
Higher expression of ECD in PC cell lines and AR regulates ECD expression in 22Rv1 and VCaP cells. **(A)** WB showing expression level of ECD in indicated PC cell lines in comparison to HPV18 immortalized epithelial cells. **(B & C)** 22Rv1 and VCaP cells were cultured in steroid free conditions for 72 hours. Cells were then treated with indicated concentrations of DHT for 12 hours and immunoblotted with anti-ECD or anti-AR antibodies. β-actin was used as a loading control. **(D & E)** qRT-PCR was performed in the same samples; RNA was isolated by standard TRIzol phenol chloroform method. 18s rRNA was used for normalization. **(F & G)** Similar protocols as mentioned above were used to perform a time response with DHT. Western blotting **(F & G)** and qRT-PCR **(H & I)** were performed on cell lysates, as above. **(J & K)** 22Rv1 and VCaP cell lines were transfected with control siRNA or AR siRNA and cultured for 72 hours. Cells were then treated with indicated concentrations of DHT for 12 hours and then lysates were western blotted with anti-ECD or anti-AR antibodies. β-actin was used as a loading control in all blots. Numbers below the blots show the quantification of band intensities after normalizing with their respective loading control, β-actin in comparison with control samples using ImageJ software. Bar graphs represent mean ± SEM from three independent experiments, each done in triplicates. Student t-test was used to calculate statistical significance *** p < 0.001, ** p < 0.01.

**Fig. S2.**
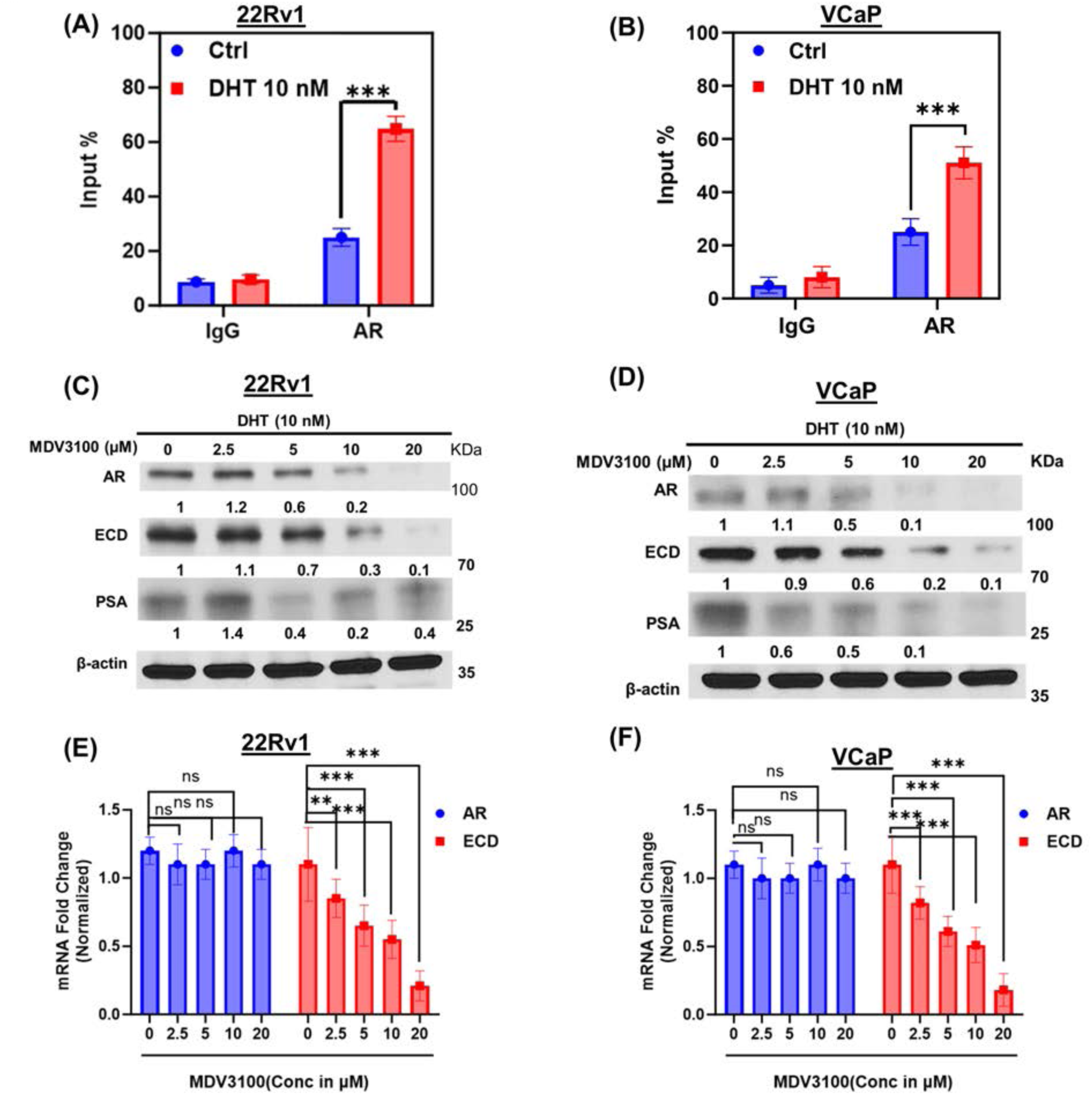
22Rv1 and VCaP cells show increased AR occupancy on ECD promoter upon androgen treatment and Enzalutamide treatment decreases ECD expression. **(A & B)** Histograms showing relative pull-down enrichment ± SEM of ECD promotor by IgG and AR from three independent experiments as observed through chromatin immunoprecipitation (ChIP) analysis. Indicated cells were cultured in steroid-free conditions for 3 days, followed by dihydrotestosterone (DHT) treatment for 3 h. Cells were cross-linked with formaldehyde and chromatin was immunoprecipitated with antibody against AR or IgG, used as a non-binding Control. Immunoprecipitated DNA was quantified with qRT-PCR for specific binding sites at ECD promoter using primers encompassing all three sites −712 to −565 from transcription start site. Student t- test was performed to calculate statistical significance *** p < 0.001. **(C & D**) 22Rv1 and VCaP cell lines were cultured in steroid-free medium for 72 hours, cells were then treated with 10 nM DHT, followed by treatment with indicated concentration of MDV3100 for another 48 hours. Lysates were collected and western blotted with indicated antibodies. β-actin was used as a loading control. **(E & F)** qRT-PCR was performed in the same samples where the RNA was isolated by standard TRIzol phenol chloroform method. 18s rRNA was used for normalization. Bar graphs represent fold change ± SEM in AR and ECD mRNA in 22Rv1 **(E)** and VCaP cells **(F)** respect to vehicle treatment from three independent experiments. *Student t- test* was performed to calculate statistical significance *** p < 0.001, ** p < 0.01.

**Fig. S3.**
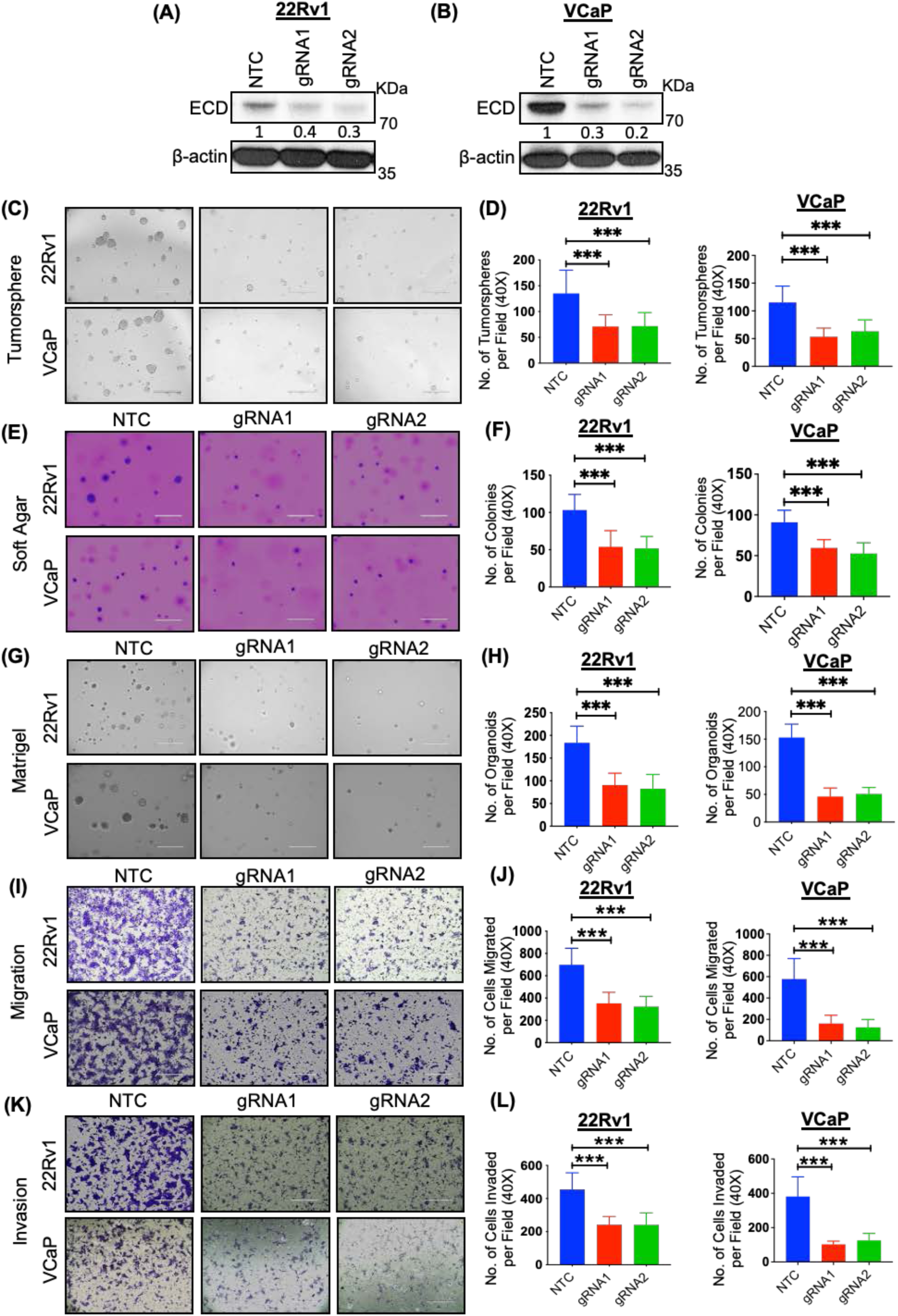
ECD depletion diminishes oncogenic traits in 22Rv1 and VCaP prostate cancer cells. 22Rv1 and VCaP cells expressing doxycycline (Dox) regulated Cas9 and guide RNAs targeting either ECD (gRNA1 & 2) or non-targeting control (NTC) were cultured in the presence of Dox (1 µg/ml) for 96 hours to deplete ECD. **(A & B)** Western blot depicting expression of ECD in 22Rv1 **(A)** and VCaP **(B)** post Dox (1 µg/ml) treatment for 96 hours. Numbers below the blot show the quantification of band intensities after normalizing with their respective loading control β-actin in comparison with NTC, using ImageJ software. Representative images and histograms depicting tumorsphere **(C & D),** soft agar colony **(E & F**), Matrigel organoid formation **(G & H),** migration **(I & J),** and invasion **(K & L),** of control and ECD depleted 22Rv1 and VCaP cells. The representative images of tumorspheres, soft agar colonies, Matrigel organoids, migration and invasion are at100 x magnification, scale bar, 400 µm. The Bar graph data presented as mean values ± SEM of three experiments, each done in triplicates. *Student t-test* was used to calculate statistical significance *** p < 0.001.

**Fig. S4.**
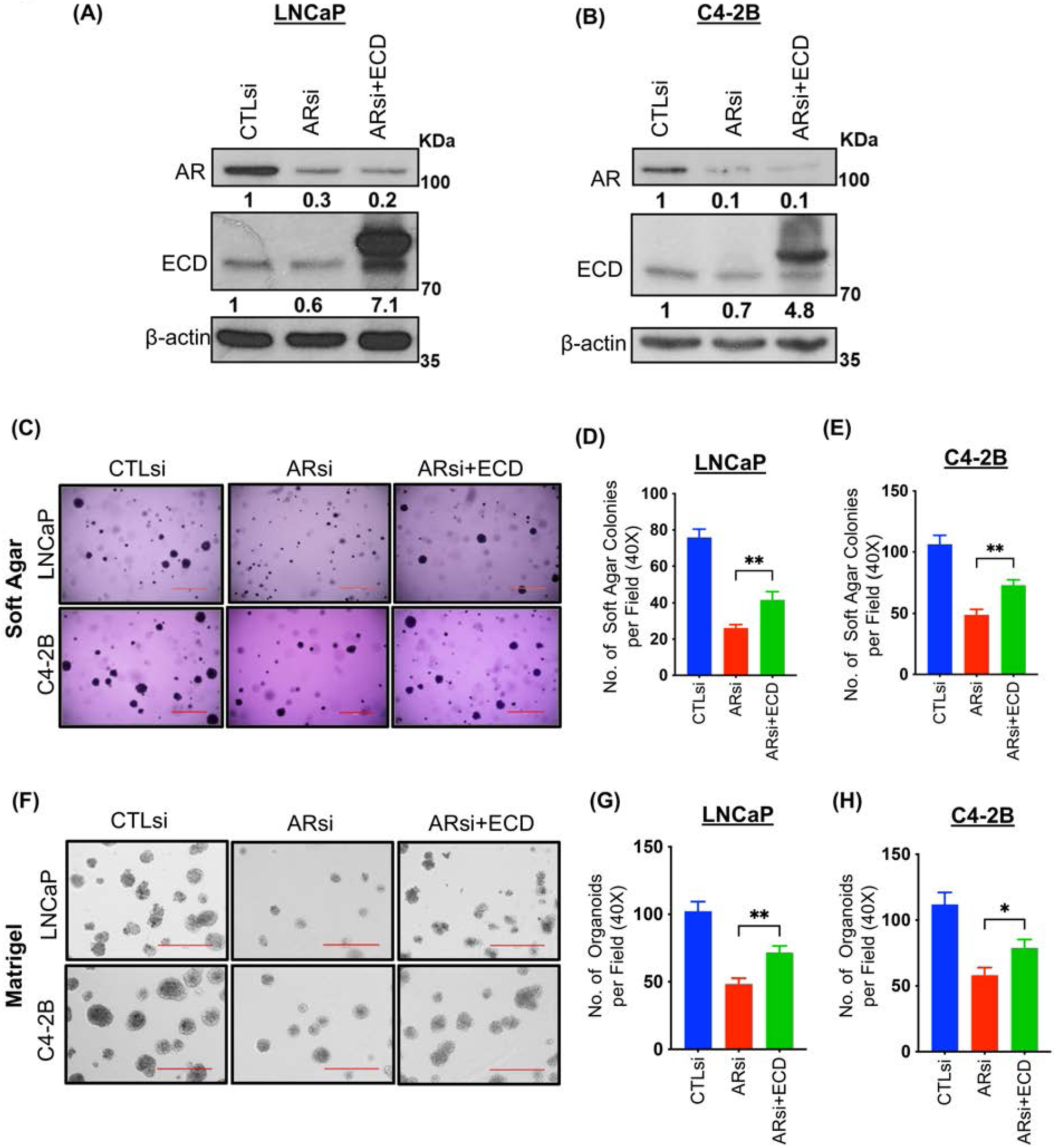
ECD overexpression partially rescues detrimental effect of AR knockdown of PC cells. LNCaP and C4-2B cells were transfected with AR siRNA and rescued with ECD overexpression. Cell lysates were collected 96 hours post transfection. **(A & B)** Western blot depicting expression of AR and ECD post AR knockdown. Numbers below the blot show the quantification of band intensities after normalizing with their respective loading control, β-actin in comparison with vector control using ImageJ software. Representative images and histograms depicting soft agar colony **(C-E**) and Matrigel organoid formation **(F-H),** of LNCaP and C4-2B post AR knockdown and ECD rescued cells. The representative images of soft agar colonies and Matrigel organoids are at 100x magnification, scale bar, 400 µm. Bar graphs show mean values ± SEM of three experiments done in triplicates from 40x magnification images. *Student t-test* was used to calculate statistical significance ** p < 0.01, * p < 0.05.

**Fig. S5.**
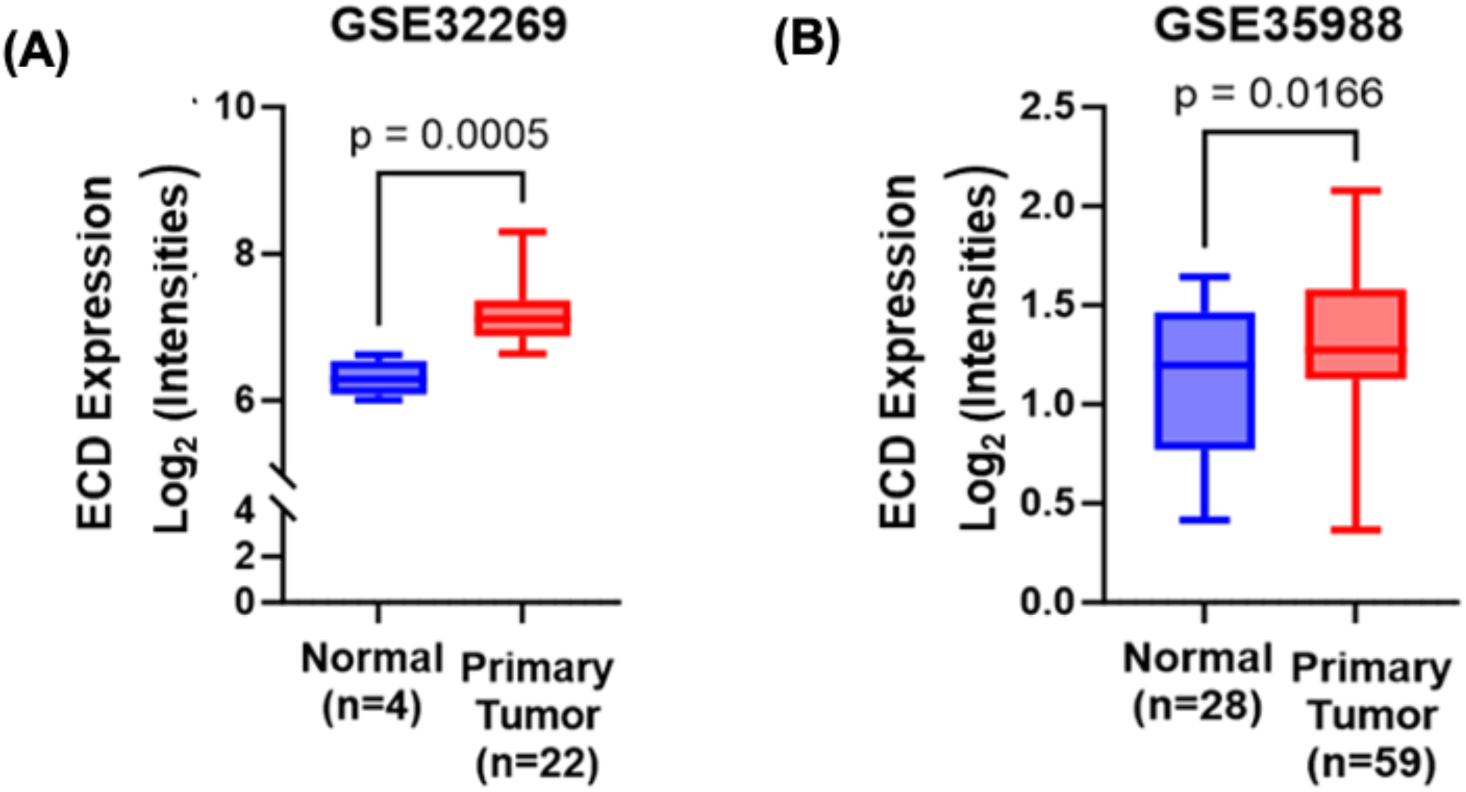
PC tissue specimens show higher expression of *ECD* mRNA. Box plot showing mRNA expression of *ECD* from **(A)** 4 normal and 22 tumor samples from cohort 1 (GSE32269) and **(B)** 28 normal and 59 tumor samples from another independent cohort (GSE35988). The mentioned two publicly available datasets were analyzed for *ECD* mRNA expression. A *paired t* test was used to compare expression in normal versus cancer samples.

**Fig. S6.**
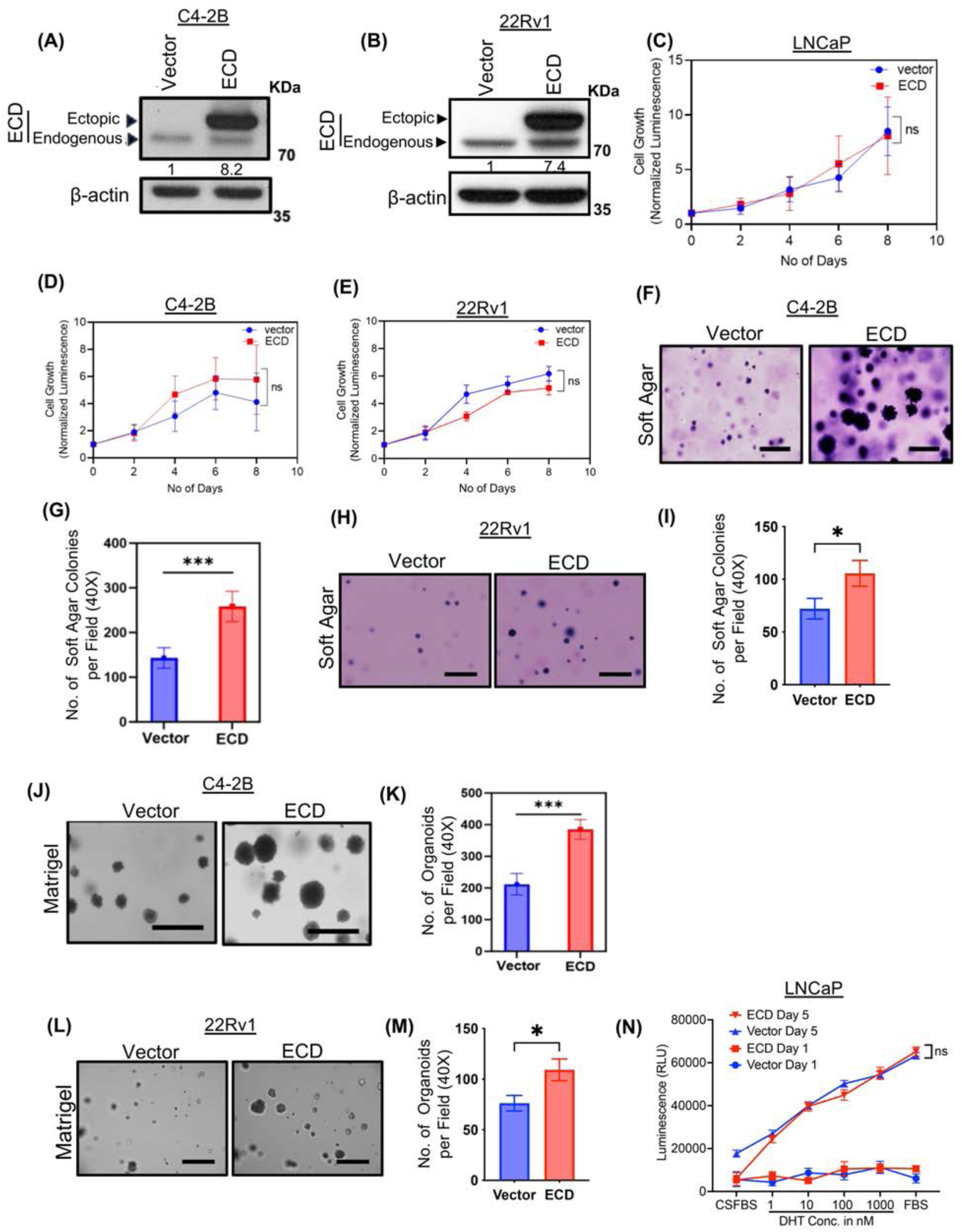
ECD-overexpressing PC cells exhibit enhanced anchorage independence and Matrigel three-dimensional (3D) growth but show no alteration in 2-D proliferation rate or Androgen dependence. **(A & B)** Western blot shows overexpression of ECD in C4-2B **(A)** and 22Rv1 **(B)** cell lines. Numbers below the blot shows the quantification of band intensities after normalizing with their respective loading control, β-actin in comparison with vector cells using ImageJ software. **(C-E)** Cell proliferation curves showing proliferation rate of LNCaP **(C),** C4-2B **(D)** and 22Rv1 **(E)** cells overexpressing ECD. Assays were performed using Cell-Titer-Glo. Luminescence values were plotted against number of days and normalized to day 1. Quantitation data represents mean ± SEM with two-way ANOVA test. *n* = 3; ns, *P* > 0.05. **(F & H)** Representative images in 100x magnification (scale bar, 400 µm) depict soft agar colony formation abilities upon overexpression of ECD in C4-2B **(F)** and 22Rv1 **(H)** cells. Images were taken after fixing and staining the colonies with 0.05% crystal violet. **(G & I)** Bars depict the mean ± SEM of soft agar colonies number per 40x magnification field in vector and ECD overexpressing C4-2B **(G)** and 22Rv1 **(I)** cells. Data represent three independent experiments, each done in triplicates. ***p < 0.001, *p < 0.05. Student t-test was used to calculate statistical significance. **(J & L)** Representative images in 100x magnification (scale bar, 400 µm) depict 3D colony formation abilities of C4-2B **(J)** and 22Rv1 **(L)** cells in Matrigel upon overexpression of ECD. **(K & M)** Bar graphs show mean ± SEM of 3D colonies number per 40x magnification field in vector and ECD overexpressing C4-2B **(K)** and 22Rv1 **(M)** cells. Data represent three independent experiments, each done in triplicates. ***p < 0.001, *p < 0.05. Student t-test was used to calculate statistical significance. **(N)** LNCaP cells expressing vector or ECD were cultured in charcoal-stripped serum containing phenol red free medium for 72 hours. Cells were then treated with indicated concentrations of DHT. Growth Assays were performed using Cell-Titer-Glo. Luminescence values were plotted for both Day 1 and Day 5. Data represent three independent experiments, each done in triplicates. ^ns^p > 0.05, Two-way ANOVA (mixed model) was used to calculate statistical significance.

**Fig. S7.**
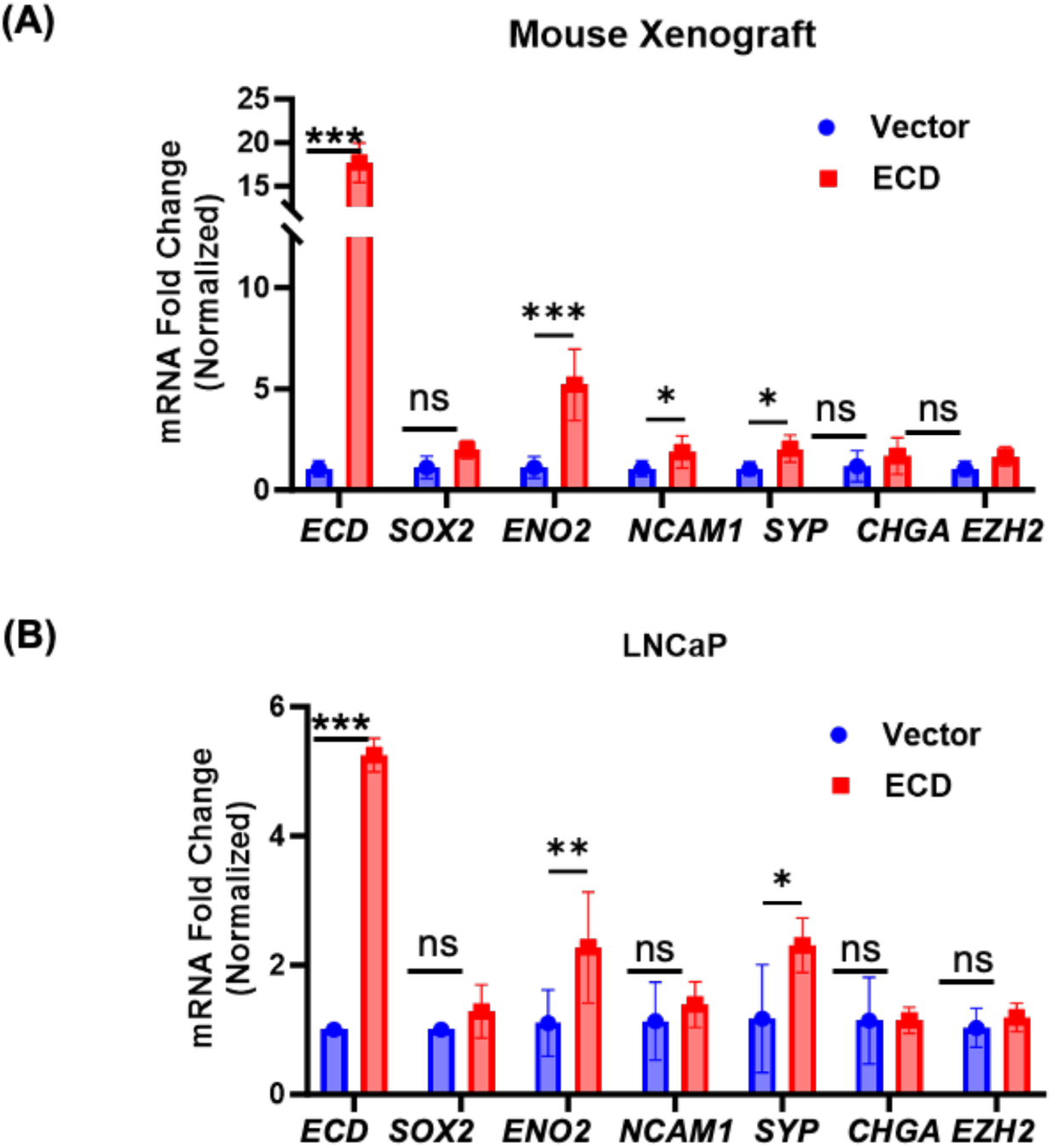
Transcriptomic analysis of ECD overexpressing prostate xenograft tumors and LNCaP cell line confirm upregulation of Neuroendocrine PC markers. **(A & B)** qRT-PCR analysis of indicated neuroendocrine (NE) signature genes of LNCaP-vector expressing and LNCaP ECD-OE xenograft tumors **(A)** and vector expressing or ECD-OE LNCaP cells **(B).** 18s rRNA was used for normalization. mRNA quantitation data represents mean ± SEM with two-tailed unpaired *t* test. *n* = 3; ns, *P* > 0.05, **, *P* < 0.01; ***, *P* < 0.001.

**Fig. S8.**
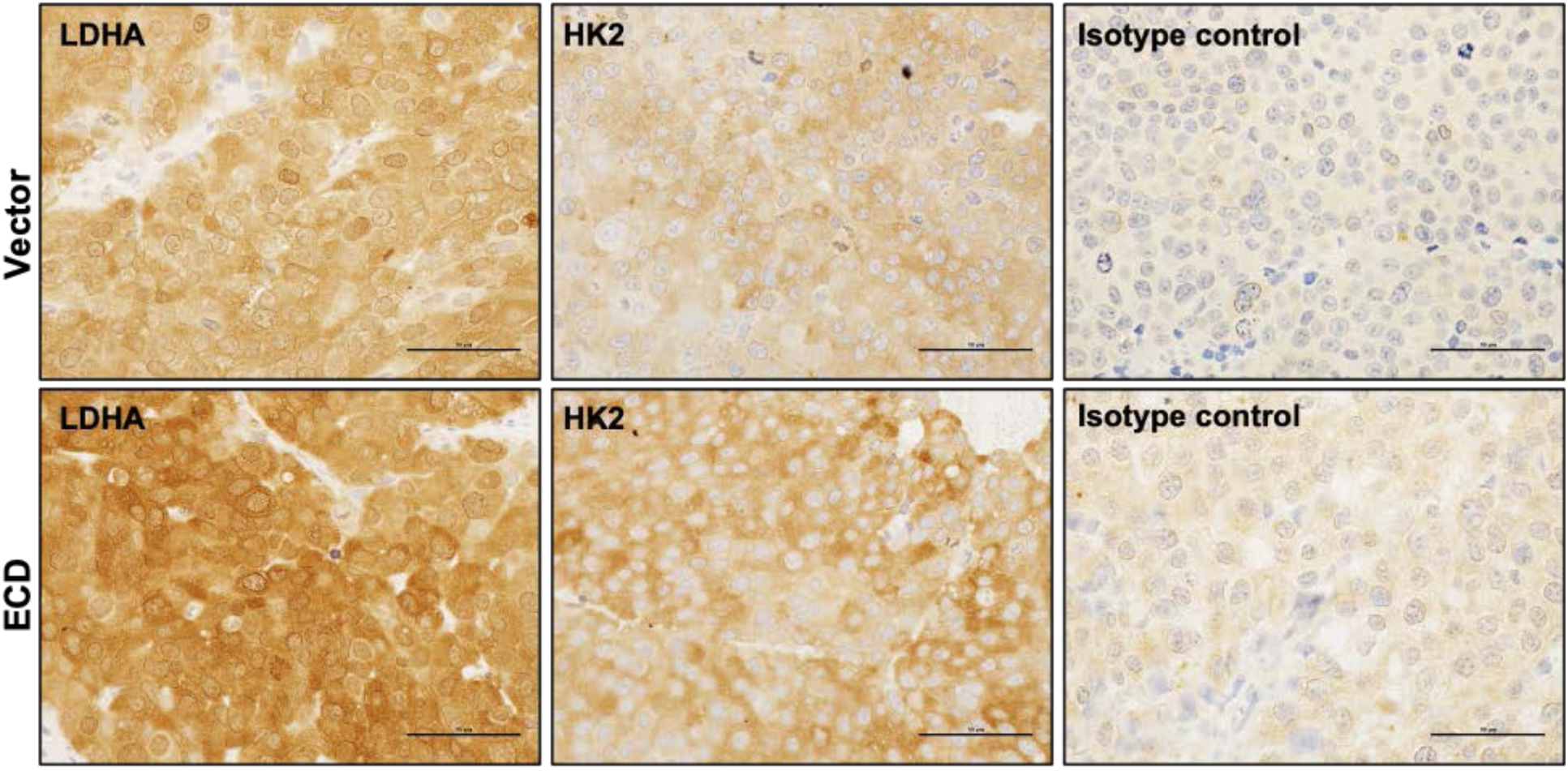
Increased expression of glycolytic enzymes, LDHA and HK2 in ECD overexpressing LNCaP tumors. IHC analysis of paraffin embedded sections of LNCaP xenografts tumors from vector group and ECD-overexpressing groups with LDHA and HK2. Rabbit IgG was used as isotype control for the staining. Representative images from both groups, in 400x magnification (scale bar, 50 µm).

**Fig. S9.**
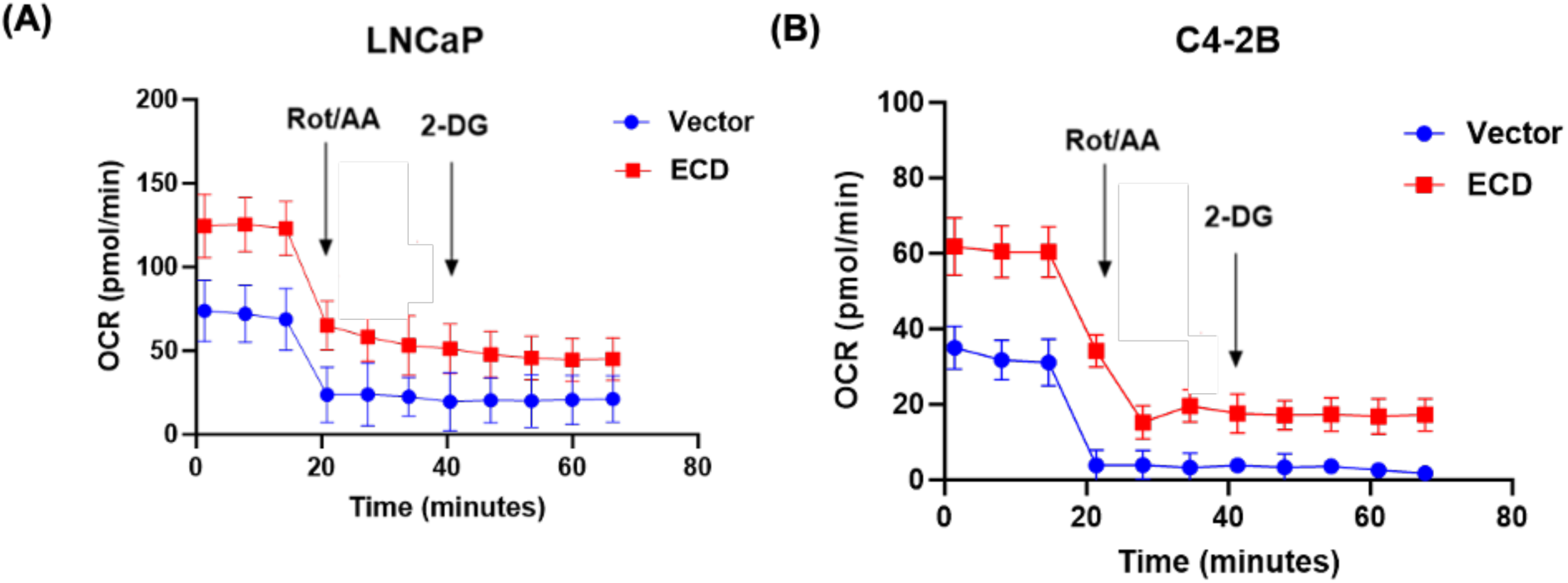
Seahorse Glycolytic Rate assay in prostate cancer cell lines overexpressing ECD. **(A & B)** Oxygen consumption rates (OCR) at various time points followed by injections of Rot/AA (0.5 μM), and 2-DG (50 mM) in vector or ECD-OE cells.

**Fig. S10.**
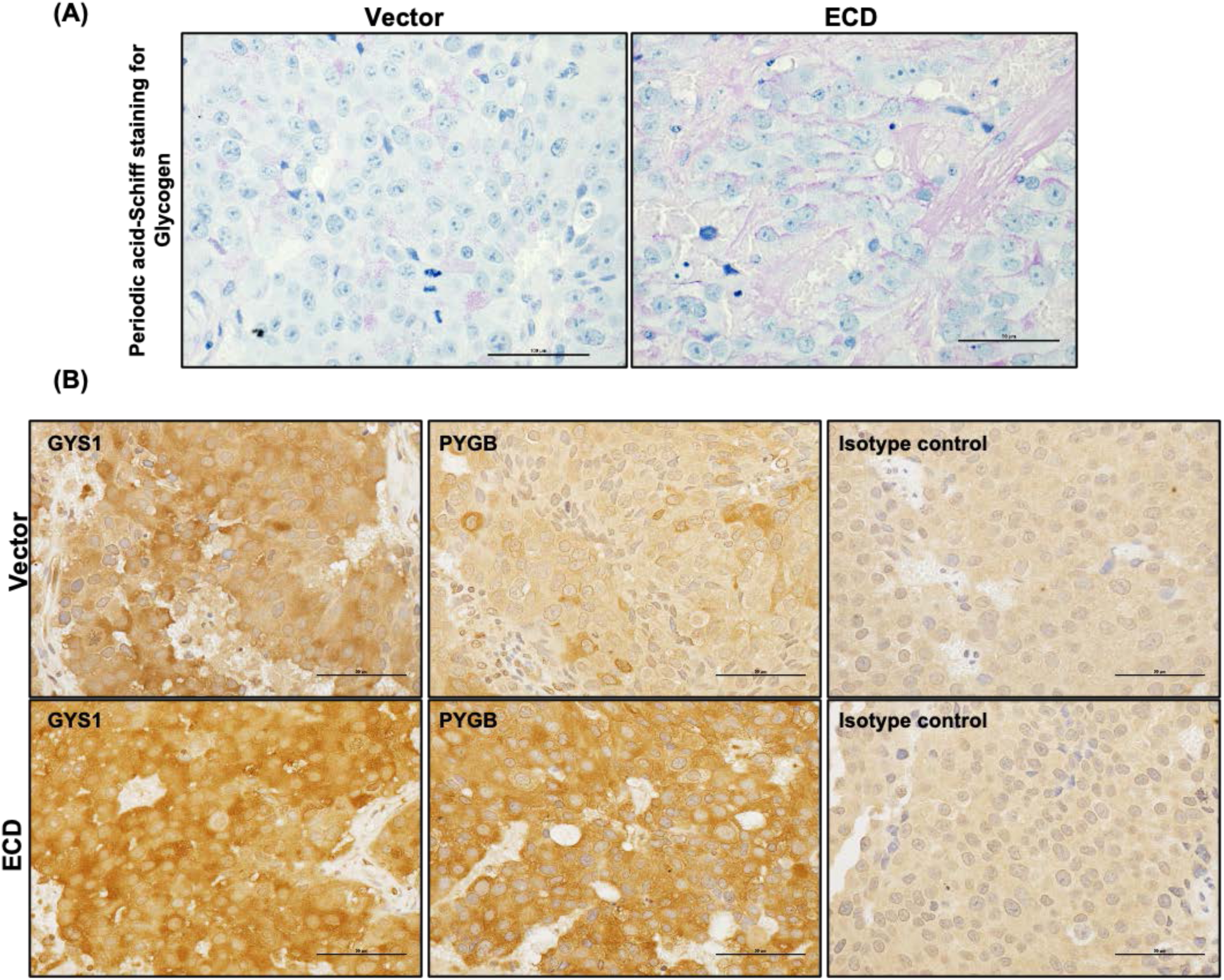
Enhanced glycogenesis and glycogen metabolism in ECD overexpressing LNCaP tumors. **(A)** Periodic acid-Schiff staining shows increased glycogen deposition represented by magenta staining in ECD overexpressing tumors compared to vector expressing tumors. (**B**) IHC analysis of paraffin embedded sections of LNCaP xenografts tumors from vector group and ECD-overexpressing groups with glycogen synthase 1 (GYS1) and glycogen phosphorylase, brain (PYGB). Rabbit IgG was used as isotype control for the staining. Representative images from both groups, in 400x magnification (scale bar, 50 µm).

